# Susceptible bacteria survive antibiotic treatment in the mammalian gastrointestinal tract without evolving resistance

**DOI:** 10.1101/2023.01.11.523617

**Authors:** Marinelle Rodrigues, Parastoo Sabaeifard, Muhammed Sadik Yildiz, Laura Coughlin, Sara Ahmed, Cassie Behrendt, Xiaoyu Wang, Marguerite Monogue, Jiwoong Kim, Shuheng Gan, Xiaowei Zhan, Laura Filkins, Noelle S. Williams, Lora V. Hooper, Andrew Y. Koh, Erdal Toprak

## Abstract

*In vitro* systems have provided great insight into the mechanisms of antibiotic resistance. Yet, *in vitro* approaches cannot reflect the full complexity of what transpires within a host. As the mammalian gut is host to trillions of resident bacteria and thus a potential breeding ground for antibiotic resistance, we sought to better understand how gut bacteria respond to antibiotic treatment *in vivo*. Here, we colonized germ-free mice with a genetically barcoded antibiotic pan-susceptible *Escherichia coli* clinical isolate and then administered the antibiotic cefepime via programmable subcutaneous pumps which allowed for closer emulation of human parenteral antibiotic pharmacokinetics/dynamics. After seven days of antibiotics, we were unable to culture *E. coli* from feces. We were, however, able to recover barcoded *E. coli* from harvested gastrointestinal (GI) tissue, despite high GI tract and plasma cefepime concentrations. Strikingly, these *E. coli* isolates were not resistant to cefepime but had acquired mutations – most notably in the *wbaP* gene, which encodes an enzyme required for the initiation of the synthesis of the polysaccharide capsule and lipopolysaccharide O antigen - that increased their ability to invade and survive within intestinal cells, including cultured human colonocytes. Further, these *E. coli* mutants exhibited a persister phenotype when exposed to cefepime, allowing for greater survival to pulses of cefepime treatment when compared to the wildtype strain. Our findings highlight a mechanism by which bacteria in the gastrointestinal tract can adapt to antibiotic treatment by increasing their ability to persist during antibiotic treatment and invade intestinal epithelial cells where antibiotic concentrations are substantially reduced.

## Introduction

The world is currently at the precipice of an antibiotic resistance crisis that is threatening to overwhelm healthcare systems in the next few decades^1^. Antibiotics are lauded as one of the most impactful medical discoveries and are credited with saving millions of lives ^2^. However, despite these successes, we have seen a dramatic rise in the incidence of antibiotic-resistant bacterial infections^3^. ∼2.8 million antibiotic resistant infections are reported per year in the United States resulting in over 35,000 deaths^4^.

Bacteria found in the gut of animals and humans account for a significant portion of the antibiotic-resistant infections and related deaths in the U.S. and throughout the world ^5^. For over forty years, we have known that these resistant bacteria can move easily between farm animals and humans, and also from humans to other humans in the hospital and in the community setting.^6-11^ Acquisition of these resistant bacteria can occur via the fecal-oral route for Gram-negative *Enterobacterales* bacteria (e.g. *E. coli, Klebsiella* spp.), which are the most common causes of urinary tract infections and among the most common causes of bloodstream infections in patients^6,12^. Hence, the mammalian gut, which is home to trillions of resident bacteria, can serve as an ideal host niche to study how gut bacteria adapt to antibiotic exposure.

One patient population particularly at risk for developing antibiotic resistant infections are immunocompromised individuals. Cancer and stem cell transplant (SCT) patients become severely immunocompromised as a result of their therapies and are treated with prophylactic and/or empiric antibiotic regimens for days to weeks to prevent serious infectious complications. Our group^13^ and others^14,15^ have shown that these patient populations develop higher incidences of antibiotic resistance and recurrent infections, leading to an increased rate of gastrointestinal carriage of antibiotic resistant bacteria resulting in excess fatality due to infections. Therefore, there is an urgent need for a mechanistic understanding for how antibiotic treatment affects gut microbiota and whether these changes/adaptations could impact the development of infectious complications.

Antibiotics given by interval dosing (intraperitoneal, intravenous, or orally) to mice are generally metabolized or cleared too rapidly to mimic serum/plasma levels observed in humans^16-18^. Therefore, we developed a murine model with the goal of better approximating the antibiotic pharmacokinetics/ dynamics observed in humans via the use of programmable subcutaneous pumps that allows for continuous antibiotic administration. We isolated an antibiotic pan-susceptible *E. coli* isolate from the feces of a pediatric stem cell transplant (SCT) patient prior to the receipt of any therapy and genetically barcoded this *E. coli* isolate. Germ-free mice colonized with this barcoded *E. coli* were then treated with cefepime, an antibiotic commonly used in cancer and SCT patients. The population size of the barcoded *E. coli* cells in the gastrointestinal (GI) tract of cefepime-treated mice substantially diminished over time, as expected from a pan-susceptible isolate, and we were no longer able to culture any viable *E. coli* from mouse feces. We, however, were able to recover *E. coli* from different segments of the GI tract (particularly the ileum), despite high cefepime concentrations in the GI tissue and plasma. To our surprise, none of the recovered *E. coli* isolates were resistant to cefepime. Rather, several of these strains had acquired genetic changes in the *wbaP* gene, which encodes an enzyme required for the initiation of the synthesis of the polysaccharide capsule and O antigen of lipopolysaccharide that allowed these mutants to survive in intestinal tissue. We were able to recapitulate these findings with an *in vitro* human colonocyte invasion assay using these recovered *E. coli* mutants as well as newly generated *E. coli* mutants lacking the *wbaP* gene. Together, our study reveals a potential mechanism by which a commensal gut bacteria exposed to high doses of an antibiotic can adapt by developing persistence and invading intestinal cells thereby avoiding high antibiotic concentrations, an observation that could have significant clinical implications as continuous antibiotic exposure can lead to antibiotic resistance^19^.

## Results

### Establishing a murine model to study the effect of antibiotics on bacteria in the gastrointestinal tract

Critically ill patients, such as cancer patients or patients in the intensive care unit, often receive prophylactic and/or empiric antibiotics administered intravenously for long durations (days to weeks)^20,21^. These hospitalized patients potentially serve as “breeding” grounds for the development of antibiotic resistance, as the resident bacteria are exposed to antibiotics and other drugs in the GI tract and ultimately could potentially translocate and cause infections^22-24^. As such, we sought to develop a preclinical model utilizing prolonged parenteral antibiotic administration to study the evolution and development of bacterial antibiotic resistance in the mammalian gut.

Administration of antibiotics (oral, intraperitoneal, or intravenous) via interval dosing to mice are often unable to mimic plasma levels observed in humans^16,18^. For beta-lactam antibiotics, maintaining antibiotic plasma levels in the plasma above minimum inhibitory concentrations (MIC) for 40-60% of the dosing interval is critical for maintaining bactericidal activity of the drug and ensuring optimal antibiotic efficacy in patients^25^. Cefepime, a beta-lactam antibiotic commonly administered to cancer and SCT patients^20,21^, requires a free time above the MIC (*f*T>MIC) of ≥ 60% for bactericidal activity^26^. We confirmed that a single subcutaneous (SC) dose of cefepime in conventional C57BL/6J mice rapidly decreased in the plasma after injection falling below the MIC for the *E. coli* isolate used in this study (0.1 µg/mL) (**Figure 1B**) by 4 hours as measured by liquid-chromatography-tandem mass spectrometry (LC-MS/MS). In contrast, cefepime administered by continuous SC infusion (20 mg/mL, 4 µl/hr) via an implanted subcutaneous pump (iPrecio 310R minipump) resulted in nearly constant cefepime plasma concentrations (∼ 5 µg/mL) over time (**Figure 1A-B**), ensuring the pharmacodynamic target of ≥60% *f*T > MIC was achieved given the cefepime MIC of the *E. coli* was ∼0.1 µg/mL. We could increase cefepime plasma concentrations by loading higher concentrations of cefepime into a SC pump, in a dose dependent fashion (**Figure 1C**). Of note, cefepime plasma levels (∼5 µg/mL) observed in mice with subcutaneous pumps loaded with 20 mg/mL cefepime was comparable to plasma trough levels (∼2-6 µg/mL) observed in humans receiving cefepime^27^, including cancer and stem cell transplant patients receiving cefepime for fever and neutropenia^28^ (**Figure 1C**). These data suggest that continuous SC delivery of cefepime in mice allows for more accurate approximation of human cefepime pharmacokinetics/dynamics.

**Figure 1.**
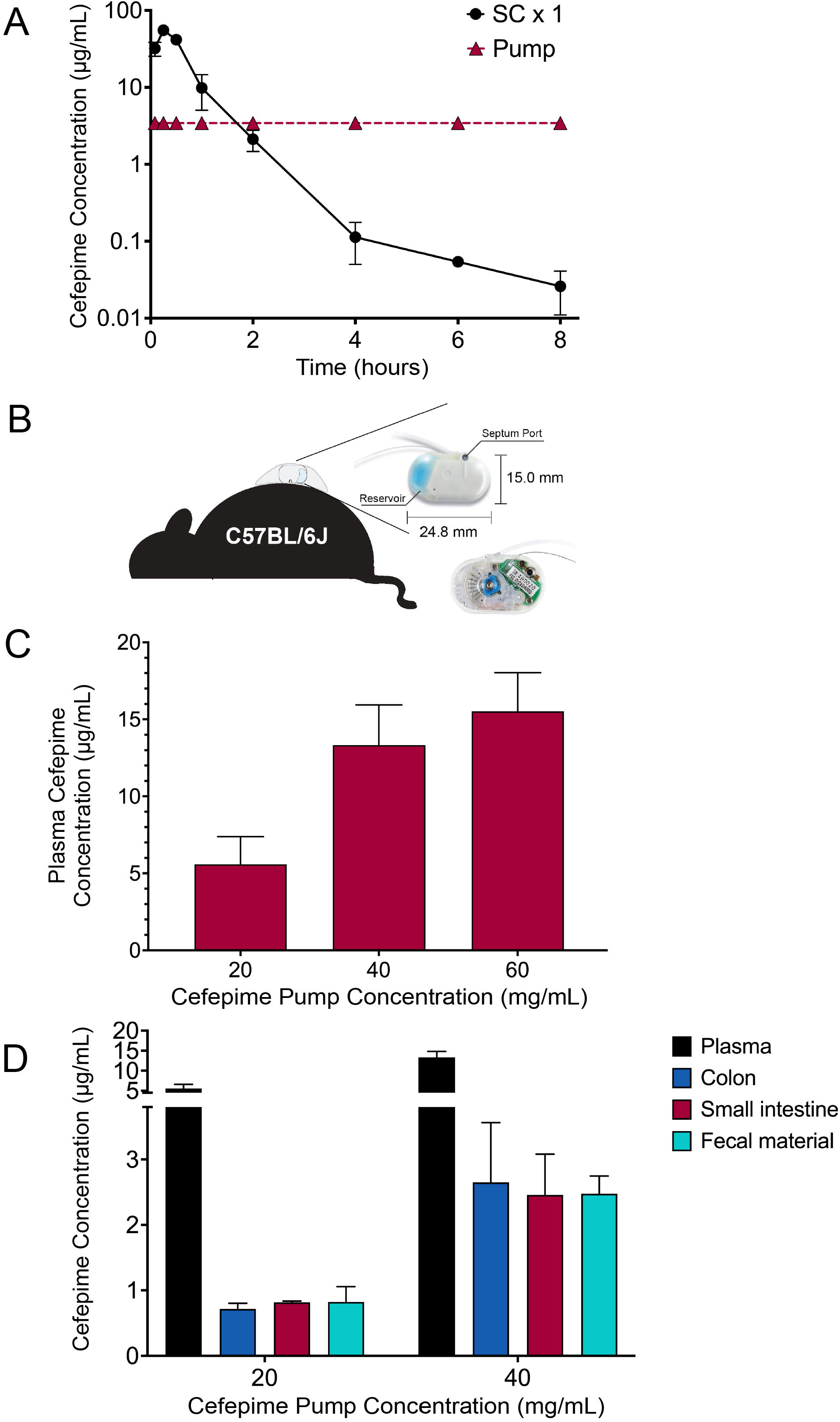
Establishing a murine model to emulate the antibiotic pharmacokinetics/dynamics observed in humans. **A**. Free cefepime concentration in plasma of mice (C57BL/6J, Jackson, female, 6-8 weeks) treated with a single subcutaneous (SC) dose (50 mg/kg dose) or via continuous cefepime infusion (iPrecio pump, 20 mg/mL, 5 µl/hr). Cefepime concentrations were calculated using Liquid-chromatography-Mass Spectrometry/MS (Shimadzu Prominence HPLC coupled to a Sciex 4000 QTRAP(r) mass spectrometer). n=3 mice per group. Points represent the mean ± SEM. Three technical replicates were performed per mouse sample. **B**. Schematic overview of subcutaneous pump (iPrecio Micro Infusion Pump System, SMP-310R, Primetech Corporation) implantation into mice. Dimensions for drug reservoir and septum (through which the drug is injected and loaded) are indicated. **C. C**. Free cefepime concentration in plasma of mice (C57BL/6, Jackson, female, 6-8 weeks) treated with continuous subcutaneous cefepime dosing (cefepime pump concentrations of 20, 40, or 60 mg/mL; @ 8 µl/hr for 20 hours). n=3 mice per group. Bars represent the mean ± SEM. Three technical replicates were performed for each mouse sample. **D**. Free cefepime concentration in plasma, colon, small intestine (Ileum) and fecal material in mice (C57BL/6, Jackson, female, 6-8 weeks) after continuous subcutaneous cefepime dosing (iPrecio pump, 8 µl/hr) for 20 hours using cefepime pump concentrations of 20 and 40 mg/mL. n=3 mice per group. Bars represent the mean ±SEM. Three technical replicates were performed for each mouse sample.

We then measured intestinal tissue cefepime concentrations in mice implanted with pumps that received cefepime (20 mg/mL) for 24 hours. Cefepime concentrations in feces, small intestine (ileum) and colon were lower than in plasma but increased with higher cefepime concentrations (40 mg/mL) loaded into the pump (**Figure 1D**). In contrast, a single oral or intravenous dose of the antibiotic levofloxacin (which is dosed once/day for both oral and intravenous administration in humans) resulted in higher concentrations in mouse intestinal tissue and fecal material compared to plasma (**Figure S1A-B**), an effect that persisted up to ∼16 hours after injection. These data are consistent with prior reports showing that parenteral administration of antibiotics result in measurable antibiotic concentration in feces in calves ^29^ and in mice^30^. In humans, oral administration of antibiotics leads to appreciable antibiotic concentrations in fecal matter^31^, but there are no comprehensive human data that we are aware of that measures corresponding plasma and intestinal tissue antibiotic concentrations after parenteral antibiotic administration.

Collectively, these data suggest that continuous subcutaneous infusion of cefepime in mice allows for a preclinical model that better mimics cefepime pharmacokinetics/dynamics observed in humans and results in measurable cefepime concentrations in intestinal tissue which could select for the development of antibiotic resistance.

### Cultivation and characterization of a pan-susceptible *E. coli* clinical isolate

To rigorously study the effect of cefepime on gut-colonizing bacteria, we isolated pan-susceptible *E. coli* strains from a fecal sample collected from a pediatric SCT patient prior to the receipt of chemotherapy and antibiotics. The rationale for choosing *E. coli* is that it is a common human gut bacterium that can readily colonize the murine GI tract and can result in disseminated bacterial infections in immunocompromised humans ^31,32^ and in preclinical mouse models^33-35^. Further, numerous *in vitro* antibiotic resistance studies have utilized *E. coli*^36-41^. Finally, as our ultimate goal is to better understand how gut bacteria adapt and evolve when exposed to antibiotics, isolating an antibiotic pan-susceptible clinical isolate recovered from human GI tract was deemed a high priority. Thus, the patient fecal sample was grown in a sequence of selective media for *E. coli* (**Figure S2A**)^42,43^. *E. coli* identification confirmation (via MALDI-TOF mass spectrometry and supplemental testing to rule-out *Shigella species*) and antimicrobial sensitivity testing by microbroth dilution was performed in the Clinical Microbiology Laboratory at Children’s Medical Center, Dallas.

An *E. coli* isolate confirmed as susceptible to all of the antibiotic compounds tested (NM43 Gram-negative MicroScan Panel, Beckman Coulter) was used for the remaining experiments in this study and was designated as PEc (<u>P</u>arental *<u>E</u>. <u>c</u>oli*). The PEc genome was sequenced (Illumina NextSeq 2000, PE 150) and the assembled genome was compared to genomes of other commonly used laboratory *E. coli* strains: the widely studied K12 substrain MG1655 ^44^; the strain BL21 commonly used for molecular cloning and recombinant protein production; and the shiga-like toxin producing O157:H7 strain which is commonly implicated in foodborne illness, though consumption of raw or undercooked foods^45^. PEc was found to have ∼100,000 base pair mismatches compared with these other *E. coli* genomes (**Figure S2B)** and thus exhibited considerable genetic distance from these other *E. coli* strains (**Figure S2C**). The contigs formed from assembling these reads were then submitted to RAST (Rapid Annotations using Subsystems Technology^46^) for functional annotation of possible genes. Notably, PEc was enriched for membrane transport proteins which consisted mainly of conjugative proteins, suggesting the acquisition of multiple mobile genetic elements when compared to the *E. coli* K12 strain BW25113 (**Figure S2D, S3**).

Next, we integrated a library of randomized DNA barcodes into the genome of the PEc, thereby creating the strain PbEc (Parental barcoded *E. coli*), to confirm the identity of and track our exogenously introduced *E. coli* clinical strain in the mouse GI tract (**Figure S4A**). Figure S4B depicts the diversity of the integrated barcodes from the initial library and the genetic distance of 20 random barcodes from each other (**Figure S4C**). PbEc did not display any detectable phenotypic deviation from the PEc, as determined by colony morphology, growth, and cefepime susceptibility (**Figure S5)**.

### Recovery of viable *E. coli* from intestinal tissue, but not fecal samples, in mice treated with cefepime

To assess the utility of our model as a tractable tool for studying antibiotic resistance, we sought to study the effect of cefepime on *E. coli* alone, in the absence of other gut microbiota. Thus, germ-free C57BL/6 mice were first colonized with PbEc via a single oral gavage. Mice then received cefepime via subcutaneous pumps (20 mg/mL cefepime, 5 µl/hr) or no antibiotics (**Figure 2A**). PbEc colonization levels in mice that did not receive antibiotic treatment, as determined by enumeration of cultured PbEc from feces and qPCR, were consistent throughout the experiment (∼10^9^-10^10^ CFU/gram feces, ∼10^5^ copies/µl) (**Figure 2B, Figure S6**). PbEc colonization levels, however, rapidly decreased and were no longer detectable in cefepime treated mice by day 6 (**Figure 2B**), even after using *E. coli* growth enrichment techniques^47^. After 7 days of cefepime administration, mice were euthanized, and intestinal tissues were harvested. Interestingly, while we were unable to culture PbEc from colonic and cecal contents (feces), we were able to isolate viable PbEc from intestinal tissue, particularly the ileum, in mice treated with cefepime (**Figure 2C**).

**Figure 2.**
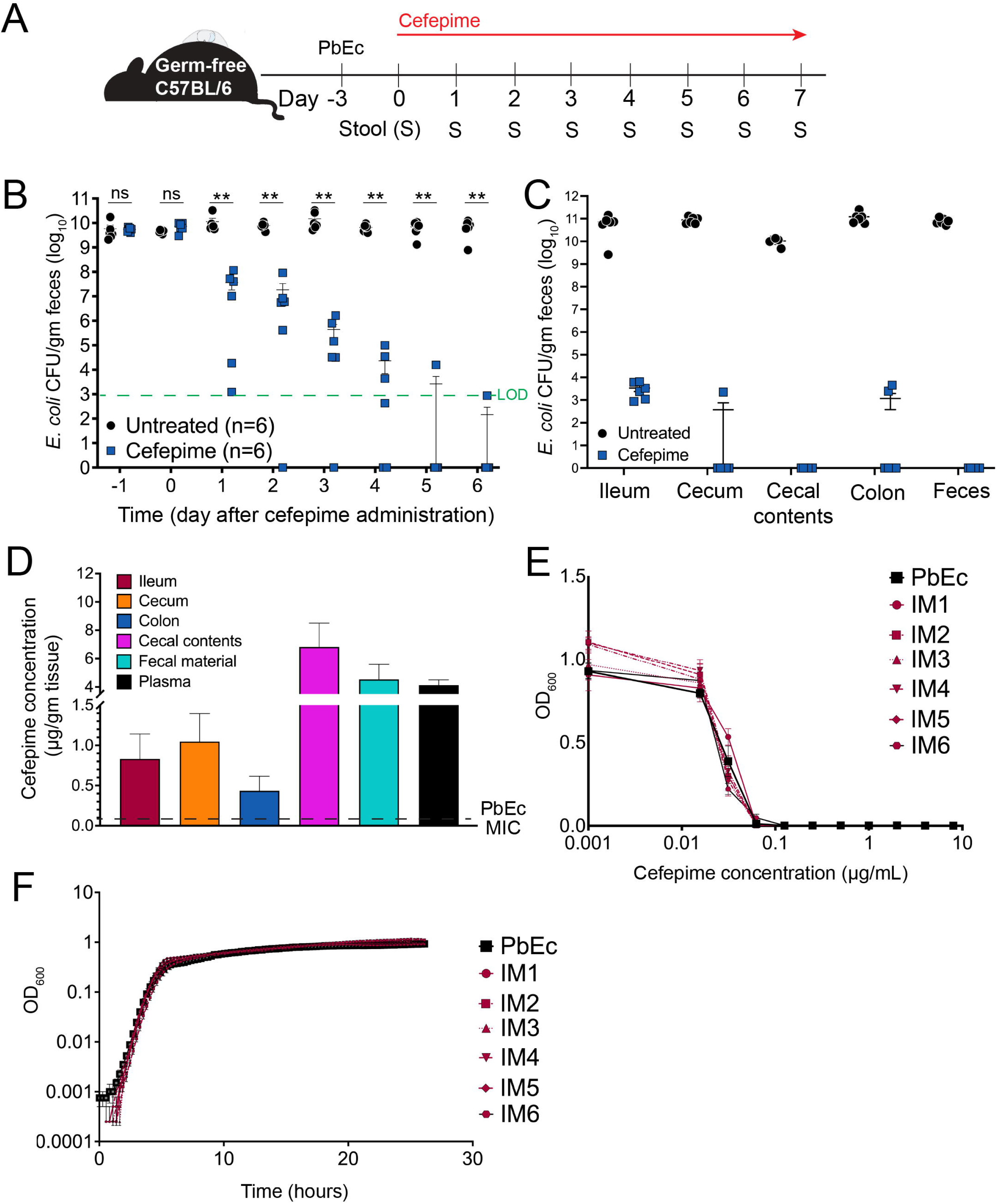
Recovery of viable *Escherichia coli* from intestinal tissue, but not fecal samples, in mice colonized with *E. coli* and treated with cefepime. **A**. Schema for germ-free mouse experiment studying the effect of cefepime on PbEc (-<u>P</u>arental <u>b</u>arcoded *<u>E</u>scherichia coli* isolate originally recovered from the feces of a pediatric stem cell transplant recipient prior to the receipt of antibiotic or chemotherapy and then genetically barcoded) colonizing the gastrointestinal tract. ***B***. *Escherichia coli* strain PbEc GI colonization levels in germ-free mice treated with or without cefepime. PbEc GI colonization levels were determined by enumeratiion of cultured *E. coli* from fecal sample homogenates. Circles (untreated) and squares (cefepime treated) represent results from individual animals. Green dashed line indicates limit of detection for the experiment. Bars represent the mean ± SEM. n=6 per group. Statistical analysis by Mann-Whitney test. *, p<0.05. ** p < 0.01. ns, not significant. ***C***. *Escherichia coli* strain PbEc levels in intestinal tissue and feces in germ-free mice treated with or without cefepime for 7 days. Intestinal tissue and fecal content were harvested, homogenized, serially diluted, and cultured on LB agar. PbEc colonization levels were then calculated by enumerating CFU and then normalizing to tissue weight. Circles (untreated) and squares (cefepime treated) represent results from individual animals. Bars represent the mean ± SEM. n=6 per group. **D**. Cefepime concentrations in intestinal tissue and feces in mice treated with cefepime for 7 days. Intestinal tissue, fecal content, and plasma was recovered from each mouse. Tissue and plasma were mixed with methanol containing 0.15% formic acid and an internal standard. Samples were vortexed, incubated at room temperature, and centrifuged. Supernatant was recovered and analyzed by LC/MS/MS to determine free cefepime concentration. Bars represent the mean ± SEM. n=6 per group. **E**. Growth of PbEc and *E. coli* isolates recovered from ileum tissue of mice treated with cefepime for 7 days (Ileal isolate from <u>M</u>ouse <u>1</u>, IM1; Ileal isolate from <u>M</u>ouse <u>2,</u> IM2, etc) in LB media with varying concentrations of cefepime. Cultures were grown at 37C with shaking (400 rpm). Growth was assessed by measuring optical density (OD_600_) after 16 hours of incubation. Two biological experiments with 4 technical replicates were performed. Points represent the mean ± SD of one biological experiment. **F**. Growth curves for PbEc and *E. coli* isolates recovered from ileum tissue of mice treated with cefepime for 7 days (Ileal isolate from <u>M</u>ouse <u>1</u>, IM1; Ileal isolate from <u>M</u>ouse <u>2,</u> IM2, etc) in LB media. *E. coli* strains were grown overnight and diluted to a final OD_600_ of 0.001. Cultures were incubated at 37°C, and growth assessed (OD_600_) every 10 minutes. Two biological experiments with 4 technical replicates were performed. Points represent the mean ± SD of one biological experiment.

Plasma and intestinal tissue cefepime concentrations in cefepime-treated mice were consistent with our prior results (**Figure 1C, Figure 2D**). Cefepime concentrations in cecal contents and feces were markedly higher than in intestinal tissues: cecal contents: 6.8 µg/g, feces: 4.5 µg/g vs ileum 0.8 µg/g, cecum 1.05 µg/g and colon 0.44 µg/g (**Figure 2D**). The survival of PbEc in illeal tissue was likely not attributable to lower cefepime concentrations, as the ileal tissue cefepime concentration (∼0.8 µg/g) was still ∼8 times higher than the MIC of PbEc for cefepime (MIC PbEc = 0.1 µg/mL). Surprisingly, there was no significant difference in either the MIC (**Figure 2E**) or the growth rate (**Figure 2F**) of PbEc isolates recovered from ileums of cefepime-treated mice (designated as Ileal isolate from mouse 1 (IM1, IM2, etc.), suggesting that PbEc survival was not a result of a canonical antibiotic resistance mechanism such as cefepime inactivation or efflux. Similarly, PbEc isolates obtained from other intestinal tissue also did not appear to have differences in MIC or growth rate **(Figure S7)**.

### Lineage tracking of barcoded *E. coli* in mice shows clonal interference and decreased population diversity during cefepime treatment

To characterize the effect of the cefepime treatment on *E. coli* populations over time, we performed *E. coli* barcode sequencing on daily fecal samples (fecal gDNA). Although hundreds of unique barcodes were detected in all mice from both treatment groups (**Figure S8**), *E. coli* populations were dominated by ∼ 30 highly abundant barcodes. Several dominant barcodes were highly abundant in most mice, possibly due to a founding effect during oral gavage of bacteria or horizontal transfer of bacteria from mouse to mouse via coprophagia. We observed highly dynamic trajectories of the 30 most abundant barcodes in both untreated and cefepime-treated populations (**Figure 3A**). These variations may have arisen due to either beneficial or costly mutations acquired by bacterial members of an asexually reproducing population in the heterogonous GI tract, a concept termed clonal interference, or horizontal transfer of bacteria between mice housed in the same cage ^48^. In the cefepime treated mice, however, some barcode lineages reached frequencies of up to ∼0.8 in the overall population, whereas this was not observed in untreated mice (**Figure 3A**). In fact, the number of unique barcodes observed in the *E. coli* populations of cefepime-treated mice was consistently lower than in the untreated control group (**Figure 3B**). This effect was most likely due to a strong population bottleneck effect where cefepime induced bacterial death greatly reduced the size and diversity of the PbEc population (**Figure 3B**), as compared to untreated mice where more lineages were prevalent throughout the experiment.

**Figure 3:**
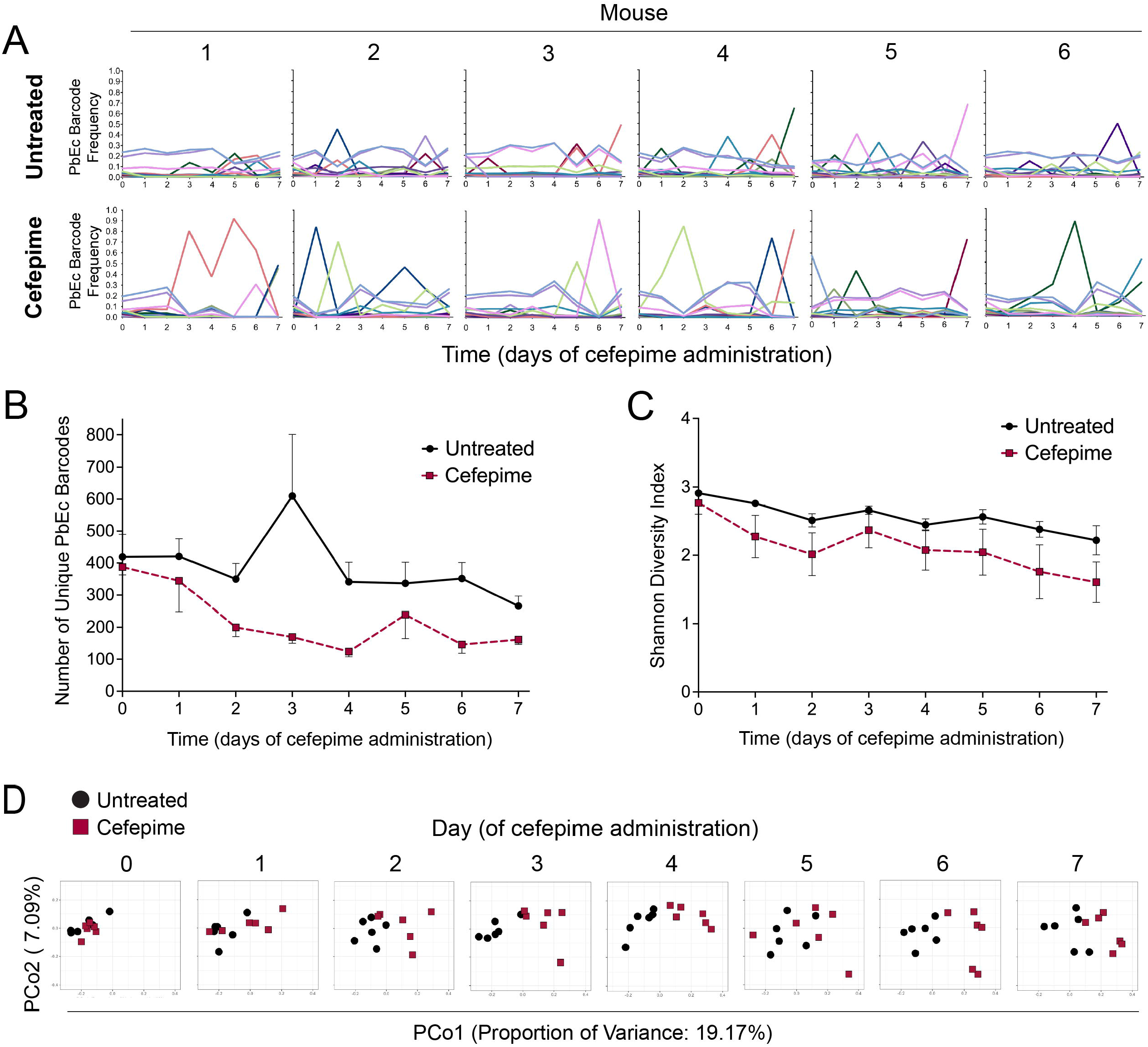
Lineage tracking of barcoded *E. coli* in mice shows clonal interference and decreased population diversity during cefepime treatment. **A**. The trajectories of the 30 most frequent barcodes in PbEc (<u>P</u>arental <u>b</u>arcoded *<u>E</u>scherichia coli* isolate) colonizing the GI tract of untreated mice (top) and cefepime treated-mice (bottom). Fecal gDNA (obtained from daily stool collection from each mouse) was isolated, and the barcode region was amplified (Primer sets: BC amp set 2 and set 3 followed by indexing using the Nextera XT DNA Library Preparation Kit). Sequencing was performed by MGIseq 2000. Reads obtained were aligned to the amplicon sequence, and barcode sequences were extracted from each sample and their frequencies were calculated accordingly. Mice were housed in four cages (3 mice per cage). Mice identified as 1, 2, and 3 were co-housed in one cage; mice identified as 4, 5, 6 were co-housed in another cage. **B**. The number of unique PbEc barcodes in mice treated with or without cefepime. Black circles (untreated) and red squares (cefepime-treated) indicate the mean number of unique PbEc barcodes in each treatment group respectively. Error bars indicate the standard deviation. **C**. PbEc barcode abundance diversity within an individual mouse (alpha diversity) was calculated using the Shannon diversity index. Black circles (untreated) and red squares (cefepime-treated) indicate the mean. Error bars indicate the standard deviation. **D**. Principal coordinate analysis was performed to assess differences in overall PbEc barcode abundance composition (beta diversity) between the untreated (black circle) and cefepime-treated (red square) groups. Treatment group dissimilarity was assessed by calculating Bray-Curtis distances (weighted and normalized). The proportion of variance accounted by each principal component is indicated.

We then assessed *E. coli* population diversity within individual mice (alpha diversity) and measured population similarity/dissimilarity between untreated and cefepime-treated groups (beta diversity). Alpha diversity (as determined by Shannon Diversity Index) declined in individual mice in both groups (**Figure 3C**), perhaps indicating that PbEc clone competition for nutrients and inhabitable niches serves as a first layer of genetic bottleneck during GI colonization. Cefepime-treated PbEc populations, however, appeared to undergo a stronger bottleneck effect, as evidenced by a greater reduction in diversity over time compared to untreated populations (**Figure 3C**). Principal component analysis (beta diversity) of PbEc barcode abundance data revealed that cefepime-treatment led to distinct patterns of PbEc population composition when compared to untreated counterparts (**Figure 3D**).

Collectively, these data suggest that lineage tracking of barcoded bacteria within the murine GI tract is a tractable tool to study the short-term and long-term effects of antibiotic exposure on bacterial population dynamics and diversity.

### *E. coli* isolated from intestinal tissue of cefepime-treated mice exhibit antibiotic persistence

To garner greater mechanistic insight into how PbEc ileal isolates were able to survive cefepime exposure, we screened bacterial isolates from individual mice for phenotypic changes. As noted above, there was no difference in either the MIC (**Figure 2E**) or the growth rate (**Figure 2F**) of PbEc isolates recovered from ileums of cefepime-treated mice compared to PbEc. However, we did notice an observable phenotypic difference (increased translucency) in several of the PbEc ileal isolates we recovered from cefepime treated mice (**Figure 4A**). Hence, we performed whole genome sequencing of PbEc ileal strains isolated from four different mice (**Figure 4B**). Two isolates -- Ileum isolate recovered from <u>m</u>ouse <u>5</u> (IM5) and Ileum isolate from mouse 6 (IM6) -- had acquired mutations in the *wbaP* gene (**Figure 4B**) which catalyzes the first step in capsular polysaccharide synthesis in *E. coli* and *Klebsiella* ^49,50^ and has also been implicated in lipopolysaccharide (LPS) O-antigen synthesis in *Salmonella* and *Porphyromonas*.^51,52^ Two different loss-of-function mutations in *wbaP* occurred independently in two different mice suggesting that loss of this gene may have had some beneficial effect for survival of *E. coli* in cefepime treated mice. Interestingly, *wbaP* capsular mutants have previously been reported to exhibit a more translucent colony phenotype compared to the wildtype strain in *Klebsiella pneumoniae*.^50,53^

**Figure 4.**
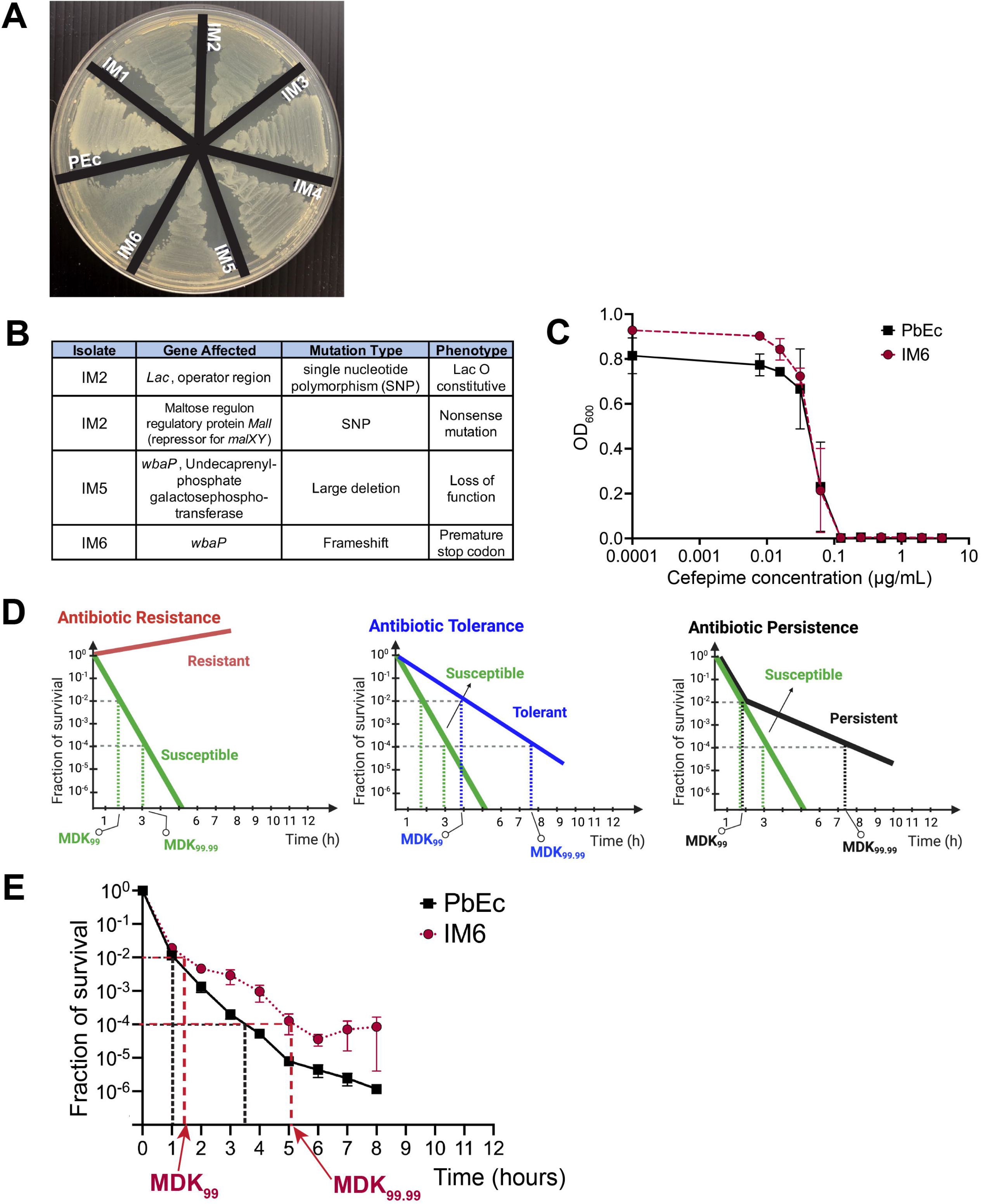
*E. coli* recovered from intestinal tissue of cefepime-treated mice exhibit antibiotic persistence. **A**. Image of select *E. coli* isolates recovered from the ileum of mice treated with cefepime (IM1-IM6) and parent barcoded strain (PbEc) grown on LB agar plate. Streaked inoculum normalized for each isolate. **B**. Table summarizing genetic mutations, mutation type, and predicted phenotypes in *E. coli* isolates recovered from the ileum in mice treated with cefepime. **C**. PbEc and IM6 growth curves in response to varying concentrations of cefepime. Cultures of IM6 and PbEc were grown overnight in LB and added to a 96-well plate containing serial dilutions of cefepime (2-fold) to a starting OD of 0.001. Plates were incubated (400 rpm, 37°C) overnight and growth assessed by OD_600_ measurement. Points represent mean ± SD. 3 replicates were performed for each experiment. **D**. Schematic figures showing illustrative defintions of hypothetical time-kill curves of bacteria exposed to antibiotics with different phenotypic outcomes: Resistant and Susceptible strains (**left**), Tolerant and Susceptible strains (**center**) and Persistent and Susceptible strains (**right**). Minimum duration for killing (MDK) values for killing 99% (MDK_99_) and 99.99% (MDK_99.99_) of total bacteria is indicated for each outcome. **E**. Time-kill assay of PbEc and IM6 after exposure to cefepime (1 µg/mL). Overnight cultures of PbEc and IM6 (n = 6 replicates) were diluted 1:1000 in LB and re-incubated at 37°C with shaking until at log phase growth. Initial timepoint (T_0_) *E. coli* quantification was performed (by enumeration of cultured CFU) and then cefepime (1 µg/mL) was added to each of the cultures and incubated at 37°C with shaking. Every hour, *E. coli* growth/survival was assessed by enumeration of cultured CFU. Points represent the mean ± SEM. Black vertical dashed lines and red vertical dashed lines are used to represent the calculated MDK_99_ and MDK_99.99_ values for PbEc, and IM6, respectively.

Because the *wbaP* gene mutation was observed in 2 of the 4 isolate sequences, we hypothesized that changes to the PEc polysaccharide capsule and/or LPS O-antigen might play an important role in the ability of PbEc to better survive antibiotic/antimicrobial peptide exposure given prior published reports^54-56^. Of note, analysis of the PEc whole genome sequencing data suggested a Group 1 polysaccharide capsule type for the PEc strain (**Figure S9A**)^57^. We could not ascertain any differences in LPS O-antigen between the ileal isolates and PbEc, as determined by performing gel electrophoresis of purified LPS from these strains (**Figure S9C**). As IM6 retained its C-terminal domain, the O-antigen synthesizing glycosyltransferase activity presumably remained intact in this strain as opposed to IM5 where the entire gene was deleted. Therefore, we proceeded to focus on strain IM6 that had a one nucleotide deletion in *wbaP*, rather than IM5 which exhibited a deletion of *wbaP*, including other genes in its operon.

PbEc isolates recovered from cefepime treated mice were not antibiotic resistant, as we found no significant difference in cefepime MIC (MIC PbEc = 0.1 µg/mL) when comparing PbEc to IM6 (**Figure 4C**). Therefore, we assessed whether IM6 exhibited an antibiotic tolerant or persistent phenotype, which is highly prevalent amongst clinical isolates recovered from patients with previous exposure to antibiotics^58-60^. Antibiotic tolerance refers to a bacterial population that can tolerate exposure to particular antibiotics, thus dying at a slower rate compared to antibiotic susceptible bacteria. Antibiotic persistence is the phenomenon by which a sub-population of cells consistently survive lethal doses of antibiotic treatment.^61-63^ Thus, adhering to the guideline definitions proposed by a multinational scientific consortium,^61^ we performed a cefepime induced time-kill assay by exposing growing cultures of PbEc and IM6 to a cefepime concentration of 1 µg/mL, which corresponds to ten times above the MIC (0.1 µg/mL). Antibiotic resistant bacteria clinically exhibit growth in the presence of antibiotic concentrations exceeding MIC, which divide antibiotic susceptibility testing into categories that correlate with the probability of clinical outcomes (illustrative definition in **Figure 4D, red line, left panel**). In contrast, antibiotic tolerant bacteria have longer minimum duration of killing for both 99 percent (MDK_99_) and 99.99 (MDK_99.99_) percent killing (illustrative definition in **Figure 4D, blue line, middle panel**), compared to antibiotic susceptible bacteria (**Figure 4D, green line, right panel**). While the MDK_99_ value for antibiotic persistent bacteria is similar to antibiotic susceptible bacteria, the time required to kill 99.9 or 99.99 percent of (MDK_99.9_ or MDK_99.99_) of persister bacteria is significantly longer (illustrative definition in **Figure 4D, black line, right panel**). In our cefepime time-kill assay, we observed that although the MDK_99_ (minimum duration for killing) was similar for both PbEc and IM6 (1.05 hrs and 1.37 hrs respectively), the MDK_99.99_ was significantly longer for IM6 (PbEc: 3.38 hrs vs IM6: 4.79 hrs, p value = 0.002, paired t-test, **Figure 4E**). Thus, we concluded that the IM6 isolates had developed an antibiotic persistence^61^ phenotype in the presence of cefepime (**Figure 4E**). Interestingly, the killing rate for both the PbEc and IM6 decreased after ∼6 hours, as has been previously described^64^. This effect is likely due to either a standing variation in both populations or degraded cefepime molecules losing their activity over time in our assay.

### *E. coli* (PEc) mutants lacking *wbaP* exhibit increased invasion of cultured human colonocytes

Bacterial capsular deletion mutants have been shown to be internalized by macrophages and bladder epithelial cells at a higher rate than wildtype counterparts.^50,65,66^ To test whether perturbations in capsule production in our clinical *E. coli* strain could play a role in colonocyte invasion, we generated a *wbaP* deletion mutant of the clinical strain PEc and performed intestinal epithelial cell invasion assays using cultured HT-29 human colonocytes^67^. We were able to culture significantly larger numbers of PEc *wbaP* deletion mutants (Δ*wbaP*) from colonocyte cells compared to colonocytes co-incubated with PEc (**Figure 5A**). Δ*wbaP* mutants complemented with *wbaP* (with arabinose-induced expression) exhibited colonocyte invasion comparable to the wildtype PEc strain. Finally, the *E. coli* IM6 isolate with a one nucleotide deletion in *wbaP* also exhibited increased invasion capacity when compared to pEc (**Figure 5B**). Interestingly, *Klebsiella wbaP* mutants have been shown to have an increased capacity to invade bladder epithelial cells, a phenomenon observed in UTI infection clinical isolates^50^.

**Figure 5.**
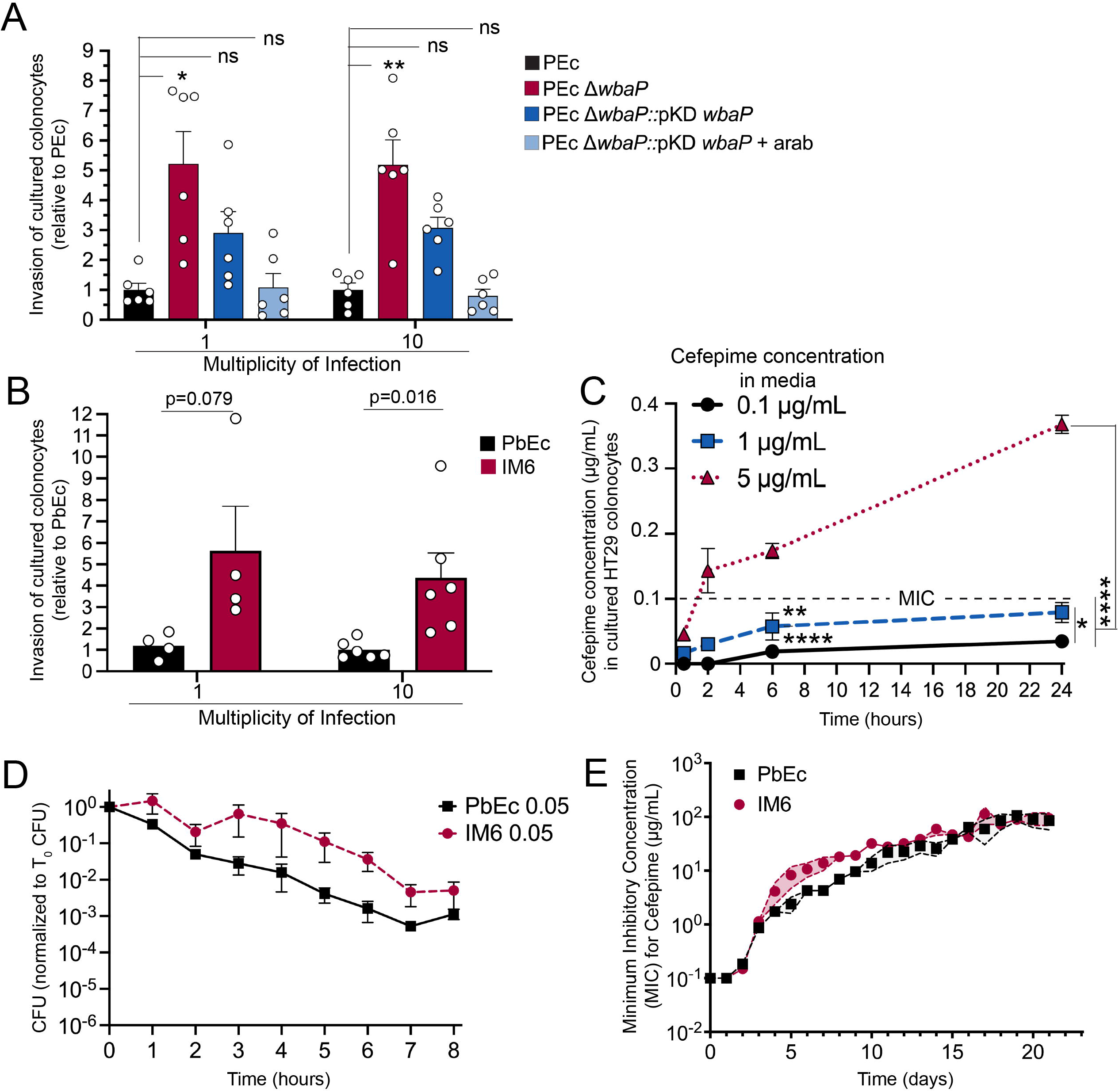
*E. coli* (PEc) mutants lacking *wbaP* exhibit increased invasion of cultured human colonocytes. **A**. Cell invasion assays of human HT-29 colonocytes using *E. coli* strains PEc (black), PEc-ΔwbaP (red), PEc-ΔwbaP + pKD-wbaP (dark blue), and PEc-ΔwbaP + pKD-wbaP + arabinose (light blue). Confluent HT-29 cells (10^5^ cells/ well) were infected with *E. coli* at a multiplicity of infection of 1 and 10 for 4 hours. Colonocytes were washed and treated with gentamicin to kill any extracellular bacteria. Colonocytes were lysed, and intracellular *E. coli* was quantified by enumeration of cultured CFU and normalized to the initial inoculum (CFU) of PEc. Points represent a single biological experiment (with 3 technical replicates). Bars represent the mean ± SEM. *, p ≤ 0.05, **, p ≤ 0.01, ***, p ≤ 0.001. **B**. Cell invasion assays of HT-29 cell lines with PbEc (black) and IM6 (red). Experimental design as described above in Fig. 5A with the exception that IM6 cell invasion numbers were normalized to PbEc. Points represent a single biological experiment (with 3 technical replicates). Bars represent the mean ± SEM. *, p ≤ 0.05. **, p ≤ 0.01. ***, p ≤ 0.001. **C**. HT-29 colonocyte intracellular cefepime concentrations. Confluent HT-29 cells were treated with DMEM media supplemented with 10% FBS and cefepime at concentrations of 5 µg/mL, 1 µg/mL and 0.1 µg/mL. HT-29 cells were harvested at select time points. The colonocyte intracellular concentration of cefepime was assessed by LC-MS/MS. n=3 replicates. Symbols (circle, square, and triangle) represent the mean ± SD. *, p ≤ 0.05. **, p ≤ 0.01. ***, p ≤ 0.001. **D**. Time-kill assay of PbEc vs IM6 at sub-MIC concentration of cefepime (0.05 µg/mL). Experimental procedure as described in Fig 4E. Mean ± SD plotted **E**. PbEc and IM6 evolved for cefepime resistance using an *in vitro* serial passage assay. PbEc (black square) and IM6 (red circle) were grown in increasing concentrations of cefepime in LB (corresponding to 0.25X, 0.5X, 1X, 2X and 4X MIC) daily. Cefepime MIC was recalculated daily, and new concentrations of cefepime were made corresponding to the new MIC. Cells grown at the highest cefepime concentration were collected and further propagated in the presence of the aforementioned range of cefepime doses. The experiment was continued for 21 days. n=6 for each group. Symbols (black square, PbEc, and red circle, IM6) represent the mean with accompanying 95 percent confidence interval values shown.

To determine whether bacterial internalization within epithelial cells provides an additional level of protection to bacteria from cefepime treatment, we measured cefepime levels within colonocytes. Cultured colonocytes were exposed to varying concentrations of cefepime added to the cell culture media: 0.1 µg/mL (MIC of PEc), 1 µg/mL (approximating our observed mouse ileal, 0.8 µg/gm, and cecal tissue cefepime concentration, 1.05 µg/gm, **Figure 2D**), and 5 µg/mL (approximating the plasma concentration of cefepime observed in our model, **Figure 1D, 2D**). Intracellular colonocyte cefepime concentration was nearly twenty times lower and directly proportional to the concentration of cefepime in the cell culture media, notably increasing over time with the highest concentration of 5 µg/mL (**Figure 5C**). Interestingly, when using a cefepime concentration of 1 µg/mL in the cell culture media, which approximates intestinal tissue levels in mice, we found that the colonocyte intracellular concentration of cefepime was below the MIC, or sub-MIC, (0.05 µg/mL) (**Figure 5C**). We next performed a time-kill assay to determine the effects of sub-MIC (0.05 µg/mL) concentrations of cefepime on PEc and IM6. At this concentration, cefepime is expected to have a significant inhibitory effect on bacterial cells but will not have a bactericidal effect as strong as what we observed at concentrations higher than MIC (e.g. 1 µg/mL, Figure 4E). We found that at sub-MIC concentrations (0.05 µg/mL), survival of both PEc and IM6 was increased approximately 10-fold, with IM6 still exhibiting higher survival compared to Pec (**Figure 5D, Figure S10**) evidenced by slower killing rate for IM6 at this lower cefepime dose. Thus, we concluded that IM6 had two fitness advantages over PEc. First, IM6 exhibits increased ability to invade cells due to the *wbaP* mutation, sheltering mutant bacteria from higher and lethal extracellular cefepime concentrations. Second, IM6 strain exhibited cefepime persistence, allowing for increased survival in both the intracellular and extracellular environment (**Figure S10**).

Finally, we tested the ability of PbEc and IM6 to evolve resistance to cefepime using an *in vitro* evolution assay. Briefly, PbEc and IM6 were passaged in varying concentrations of cefepime (corresponding to 0, 1/4X, 1/2X, 1X, 2X and 4X the MIC of PbEc) over 21 days. MIC’s were evaluated daily, and cefepime concentrations in evolution experiments were adjusted accordingly. Cells were then passaged in the aforementioned concentrations (calculated using the new daily MIC value). We found that IM6 evolved resistance at a comparable rate to PbEc (**Figure 5E**) and acquired similar mutations **(Supplementary Table 2**), suggesting that IM6 was neither more nor less adept at evolving towards cefepime resistance. Populations derived from evolving both PbEc and IM6 under cefepime selection *in vitro* had steadily increasing number of surviving *E. coli* (**Figure S11**) as conceptually described in Figure 4D. Interestingly, mutations in the *wbaP* gene were observed in evolving PbEc populations. Frequent mutations in the *wbaP* gene were previously in *K. pneumoniae*, particularly when bacterial cultures were passaged in rich growth media under aerobic conditions^68^. Previous studies have shown that repeated exposure to sub-MIC doses of antibiotics increases the propensity of bacteria to develop resistance^69-71^. Therefore, consistent exposure to sub-MIC conditions (such as those observed here) over a longer period of time could potentially lead to eventual resistance development in clinical settings.

## Discussion

Studies on antibiotic resistance development and evolution are often limited by the use of well-controlled systems utilizing permissive, and perhaps non-physiologic, conditions. And thus the conclusions arrived at using these approaches may lack clinical relevance^72^. Here, we describe a novel *in vivo* model for studying the effects of antibiotic treatment on *E. coli* residing in the gastrointestinal tract. We sought to emulate exposures observed in human patients by first focusing on achieving clinically relevant trough plasma levels. We found that antibiotic concentrations varied significantly in different intestinal tissues from the measured plasma levels, leading to recovery of bacterial isolates from only certain GI tract segments, highlighting the significance of tissue-specific antibiotic concentrations.

The mechanisms of bacterial survival after antibiotic treatment are poorly understood. Intracellular invasion has been widely implicated in facilitating these infections particularly in the case of recurrent UTI infections^73,74^: intracellular bacteria can escape membrane-bound phagocytosed vacuoles to establish cytosolic intracellular bacterial communities that can spawn reinfection and induce lower urinary tract symptoms with no apparent bacteriuria. The *wbaP* mutant, IM6 isolated from the intestinal tissue of the cefepime treated mice, as well as the *E. coli wbaP* mutant that we generated for this study, exhibited increased capacity to invade cultured human colonocytes. This phenotype has been observed previously in hypocapsulated *Salmonella enteritidis* mutants, which although previously implicated in their enhanced ability to invade macrophage cells, were also found to be defective in swarm motility which is important for proliferation in the intracellular environment ^66^. It is possible that the intracellular sub-MIC concentration of cefepime would not allow for the formation of intracellular bacterial communities or even for the bacteria to escape their initial phagocytosed membrane-bound organelles remaining in a quiescent state within these vesicles, but further investigation is required to rigorously address these questions.

Understanding and potentially predicting antibiotic resistance evolution should be an important consideration when administering antibiotics to human patients. Currently empiric antibiotic therapy is the paradigm when administering antibiotics to populations at high risk for developing infections (e.g. fever and neutropenia for cancer patients, presumed sepsis in ICU patients, etc.) with screening performed afterward to confirm the susceptibility of the infectious agent^75^. Alternatively, by understanding potential pathways to resistance, more informed and effective treatments could be prescribed that minimize the potential for resistance or cross-resistance and maximizes positive treatment outcomes. Thus, our preclinical model has the potential to generate potentially impactful insights into how microorganisms adapt and evolve when exposed to antibiotics.

Our study also highlights the utility of this model to study bacterial population dynamics and evolution. Here, we chose to study the effect of a single antibiotic on a single gut bacterium. Any genetically barcoded microorganism could be introduced into a germ-free or conventional GI tract. Potentially, any antibiotic or combinations of antibiotics could be administered via subcutaneous pumps and conventional delivery systems. Any number of additional “stressors” or variables could then be added as well: changes in diet, co-infection with a microbial pathogen, induction of an autoimmune phenotype, etc. In addition to ascertaining the effect of these stressors on the microorganism in question, the effects on the host – ranging from effect on the microbiome, metabolome, immune function, and host metabolism – could easily be interrogated as well.

In summary, we have developed a tractable and clinically relevant preclinical model to study the effects of antibiotics on bacteria residing within the gastrointestinal tract. In this study, we documented one mechanism by which commensal gut bacteria can adapt to antibiotic exposure by invading intestinal cells and exhibiting increased antibiotic persistence. In the future, this model could easily be adapted to study numerous aspects of microbiology, infectious diseases, ecology, and evolution.

### Limitations of the Study

*E. coli* colonizing the GI tract of a germ-free mouse will likely behave quite distinctly from *E. coli* colonizing the GI tract of a mouse or human with an intact and mature microbiome. Therefore, future studies will involve introducing the PbEc into conventional mice with intact gut microbiomes and then administering antibiotics. Further, as it is well documented that different inbred mouse strains harbor different gut microbiomes (and even the same strain, such as C57BL/6 mice, from different vendors have unique gut microbiomes ^76^), it would be important to confirm whether our findings are generalizable in other inbred strains of mice.

We are cognizant that the antibiotic trough levels reported in human patients arrive through bolus dosing at regular intervals. Further, we realize that bolus dosing itself, as there is an accompanying peak level prior to reaching a sustained trough level, could have significant impacts on antibiotic PK/PD observed systemically and in the gut. As such, we plan to implement bolus dosing in addition to continuous antibiotic administration in future versions of this model as we strive to better approximate the human condition (Figure S1).

### Quantification and statistical analysis

All data sets were tested for normality (e.g. Shapiro-Wilk). Data sets with normal distribution will be analyzed by parametric tests, including Student’s T-test or one- or two-way ANOVA. Non-parametric tests, such as Mann–Whitney U test or Kruskal-Wallis, were used for non-normal distributions

## Author Contributions

M.R., E.T., P.S., N. S. W., M.L.M. and A.Y.K, designed the research; M.R., P.S., L. A. C., S. A., C. B., and X. W. performed experiments. M.R., M.S.Y., J. K., S.G., N. S. W., E.T. and A.Y.K. analyzed data. M.R., E.T. and A.Y.K wrote the manuscript. All authors revised the manuscript and approved its final version.

## Materials and Methods

### Bacterial strains and growth conditions

*Escherichia coli* cultures were routinely grown in Lysogenic Broth (LB) medium aerobically (shaking at 230 rpm) at 37°C, unless otherwise specified. Growth was determined by spectrometry (OD_600_) or enumeration of cultured colony forming units (CFU) on LB plates. When specified, kanamycin was supplemented at 50 µg/mL, chloramphenicol was supplemented at 25 µg /mL, and ampicillin was supplemented at 100 µg /mL.

### Polymerase Chain Reaction (PCR)

Routine PCR was performed using Q5 High-fidelity Mastermix (NEB M0492L). PCR purification and gel extraction was performed using Macherey-Nagel Kit cat. 740609, plasmid purification was performed using Macherey-Nagel Kit cat. 740588, gDNA was extracted using QIagen DNeasy Blood and Tissue kit (cat. 69504) unless otherwise specified.

### Plasmids used

An integration cassette containing two regions of homology flanking a kanamycin-resistance gene was engineered on a pUC backbone plasmid encoding ampicillin resistance using the NEBuilder HiFi DNA Assembly mastermix (NEB E2621L). A counter-selectable marker based on PheS^77^ was engineered under the constitutive SpeI promoter and cloned into the pUC plasmid (generating pUC-BC) as well as the lambda recombineering plasmid pKD46. In addition, the ampicillin resistance marker on pKD46 was replaced by a chloramphenicol resistance cassette yielding plasmid pKD-PC. Phosphorylated primers containing one half of the barcode each at the 5’ ends were used to linearize the pUC-BC plasmid and the resulting amplicon was ligated using T4 DNA Ligase (NEB M2200L). The purified ligation reaction was transformed into NEB-10-Beta competent cells (NEB C3020K) and the lawns resulting from this transformation were pooled together and inoculated into LB broth supplemented with kanamycin. Plasmid purification from the library was performed using the Qiagen Plasmid Midi Kit (Qiagen 12123).

### Barcoding protocol

The clinical strain PEc was first transformed with the modified lambda recombineering plasmid pKD-PC. The resulting transformants were induced for lambda recombination using arabinose, and the resulting cells were transformed with 1 microgram of the barcoded plasmid library. After overnight incubation at room temperature, the cells were plated on pre-warmed M9-Kanamycin-4-chloro-phenylalanine plates (200 mL 5X M9 salts, 100 mL 10X Amikacin, 40 mL 25X Glucose, 2 mL MgSO4, 100 µl CaCl2, 2 gr 4-Chloro-DL-phenylalanine (Alfa Aesar, cat. A13323), 15 gr agar per 1 L of media). After overnight growth at 40°C, the lawns of cells were scraped and diluted in M9 broth supplemented with kanamycin and 4-chloro-DL-phenylalanine. Several passages were performed taking at least 10^6^ cells per passage to completely remove ampicillin and chloramphenicol resistance genes from the library. To assess integrated barcode diversity, amplification of the barcodes was always performed using one primer outside of the region of homology on the chromosome while the other primer was located within the integration cassette on the other side of the barcode.

### *wbaP* deletion protocol

Two regions of homology corresponding to the first 100 bp and the last 100 bp of the *wbaP* gene were cloned into a pUC-origin plasmid with a chloramphenicol-resistance cassette flanked by FRT sites in between them. Constructed plasmid was sequence confirmed and subsequently amplification of the deletion construct was performed using Δ*wbaP* set 1 forward primer and Δ*wbaP* set 3 reverse primer (Supplementary Table 1). This amplicon was transformed into the PEc strain induced for lambda recombineering^78^ and plated on LB-chloramphenicol.

Colonies appearing from this transformation which appeared translucent were inoculated into LB-chloramphenicol media and incubated overnight at 40°C to cure the lambda recombineering plasmid. The following day, glycerol stocks of the deletion were made and a portion of the culture was also plated on ampicillin to ensure removal of the pKD46 plasmid.

The chloramphenicol resistance cassette was subsequently removed using aprotocol^78^ with the plasmid pCP20 targeted recombination at the FRT sites.

### *wbaP* complementation

*wbaP* complement primers listed in Supplementary Table 1 were used to create 1) a plasmid bearing pUC origin; 2) the arabinose-inducible promoter construct from pKD46, and 3) the full *wbaP* gene from PEc forming the plasmid pKD-wbaP. *wbaP* expression was induced using 50 mM L-arabinose.

### Mice

C57BL/6J (strain number 000664) mice were obtained from Jackson Laboratories and bred and maintained in the barrier facility at the University of Texas Southwestern Medical Center. All animals were kept on a 12-hour light-dark cycle and were fed standard mouse chow (Teklad 2916, irradiated). Female, 5-7 week old mice were used for all experiments. Experiments were performed using protocols approved by the Institutional Animal Care and Use Committees of the UT Southwestern Medical Center.

Germ-free C57BL/6 mice were maintained in isolators as described.^79^ A sample size of six per group was chosen using the resource equation method^80^, and as the best compromise between providing an adequate sample size for assessing differences in colonization balanced with the limited availability of germ-free mice. Inclusion/exclusion criteria were not established. No animals were excluded from analysis. Randomization was not used, and no blinding was done for mouse studies.

### Determination of cefepime dosage

#### LC-MS/MS analysis of cefepime

Levels of cefepime in plasma, tissues, and cell lysates were determined by LC-MS/MS. Briefly, plasma, tissue homogenate, or cell lysate was mixed with a two-fold volume of methanol containing 0.15% formic acid and 150 ng/mL of the internal standard ceftazidime. Samples were vortexed for 15 seconds, incubated for 10 min at RT and then centrifuged for 5 min at 4°C for 5 min at 16,100 x g. The supernatant was centrifuged a second time before placing in an HPLC vial. Levels of cefepime were monitored by LC-MS/MS using an AB Sciex (Framingham, MA) 4000 QTRAP® or Triple Quad 4500™ mass spectrometer coupled to a Shimadzu (Columbia, MD) Prominence LC. The compound was detected with the mass spectrometer in positive MRM (multiple reaction monitoring) mode by following the precursor to fragment ion transition 481.1/396.0. An Agilent C18 XDB column (5 micron, 50 × 4.6 mm) was used for chromatography with the following conditions: Buffer A: dH20 + 0.1% formic acid, Buffer B: methanol + 0.1% formic acid, 0 - 0.5 min 5% B, 0.5 – 1.5 min gradient to 100% B, 1.5 - 4.5 min 100% B, 4.5 - 4.6 min gradient to 5% B, 4.6 - 5.5 5% B, flow rate 1.5 ml/min. Ceftazidine (transition 547.101/468.0) from Sigma (St. Louis, MO) was used as an internal standard (IS).

#### Sub-cutaneous pump administration of cefepime and assessment of tissue cefepime concentrations

Conventional C57BL/6 mice (female, 5-7 weeks age) were anesthetized with isoflurane and then implanted with Alzet model 2001D mini-osmotic pumps subcutaneously. Pumps were then filled with 20, 40 and 60 mg/mL of active cefepime (PremierProRx, Sagen Pharmaceuticals) prepared in sterile HPLC grade water. Mice were administered 5 mg/kg carprofen SC at time of surgery and then 18 hours later for pain management. 20 hours after surgery, the animals were euthanized by inhalation overdose of CO_2_ and pumps were removed. Whole blood was harvested by terminal cardiac puncture. Plasma was processed from whole blood by centrifugation of the ACD treated blood for 10 minutes at 10,000 rpm in a standard centrifuge. Lung, colon, small intestine, small intestine contents, and fecal pellets were also collected. The tissues were weighed and snap frozen in liquid nitrogen. Tissues were homogenized in a three-fold volume to weight ratio of PBS using a polytron homogenizer (IKA T25 Ultra Turrax Basic Homogenizer, IKA Works, Wilmington, NC). Homogenates were stored at -80°C until analysis by LC-MS/MS.

#### Comparison of different dosing methods for cefepime Subcutaneous (SC)

C57BL/6J mice were administered 50 mg/kg cefepime SC (0.2 ml per SC injection). Whole blood was harvested by cardiac puncture from euthanized mice and placed into acid citrate dextrose (ACD) tubes. Plasma was extracted after centrifugation (9,600 x g for 10 minutes). Colon, small intestine, fecal material, and fecal pellet were also collected at this time. The tissues were weighed and snap frozen in liquid nitrogen. Tissues were homogenized in a three-fold volume to weight ratio of PBS as previously described. Homogenates were stored at -80°C until analysis by LC-MS/MS.

#### Pump only

A separate cohort of animals were implanted SC with iPrecio pumps (iPRECIO Micro Infusion Pump System, SMP-310R; Primetech Corporation, Tokyo, Japan) containing 20 mg/mL Cefepime (flow rate: 4 µL/hr) and plasma and tissue levels determined in the same manner. Exposure to the combination was simulated mathematically

#### Isolation of *Escherichia coli* isolate from patient fecal sample

After initial enrichment in lactose broth, successive plating on selective agar: mEndo followed by mFC agar and finally MacConkey Agar with MUG. Colonies appearing fluorescent on this media were identified as *E. coli* (designated PEc) using MALDI-TOF (Bruker MALDI Biotyper); genome sequencing ruled-out potential misidentification of *Shigella* species by MALDI-TOF.

#### Genomic analyses of PEc strain

Genomic DNA was isolated from the clinical strains obtained using DNeasy Blood and tissue kit (Qiagen cat. 69504). DNA was sent to Microbial Genome Sequencing Center, PA (now Seqcenter) for Illumina whole genome sequencing. Paired-end reads (2 × 150 bp) were first assembled into contigs using CLC Genomics Workbench *de novo* assembly tool. These contigs were then submitted to RAST for annotation and prediction.

Trim Galore (https://www.bioinformatics.babraham.ac.uk/projects/trim_galore/) was used for quality and adapter trimming of Illumina sequencing reads. Unicycler (v0.5.0) (Wick et al., 2017) was used to generate de novo genome assembly from Illumina and Oxford Nanopore sequencing reads. RefSeq genome assemblies of E. coli MG1655, BL21, and O157:H7 were downloaded from National Center for Biotechnology Information (NCBI) FTP site. MUMmer 4 (Kurtz et al., 2004) was used to compare the genome assemblies. The mutation rate between two genomes was calculated by the number of SNPs divided by the alignment length and Jukes-Cantor distances were calculated from the mutation rates. The neighbor-joining tree was generated based on the distances using R-packages ape (Paradis et al. 2019) and ggtree (Yu, 2020).

#### Preclinical model studying the effects of antibiotics on *E. coli* in the GI tract

5-7 week old female C57/BL6 germ-free mice were used, with n=6 for each group. Mice were gavaged with 1 ×10^7^ CFU PbEc. 2 days after oral gavage, PbEc GI colonization levels were assessed by enumeration of cultured PbEc from fecal samples. On Day 3 after oral gavage, 6 mice were implanted with iPrecio SMP-310R pumps which were filled with cefepime 20 mg/mL dispensing 5 µl/hr. Stool was collected daily for measurement of PbEC GI levels. Pumps were refilled with cefepime daily afrer mice were briefly anesthetized using isoflurane. On day 10 after oral gavage, mice were euthanized, and GI tract tissue (ileal, cecal, colon, fecal material from cecum and colon) were harvested. A portion of fresh GI tissue was homogenized and incubated in lactose broth to assess for PbEc growth. The remainder of GI tissue was snap frozen in liquid nitrogen. Tissue was later used for measuring cefepime concentrations via LC-MS/MS analysis and extracting genomic DNA for sequencing.

#### *Escherichia coli* growth quantification from mouse tissue

Stool was collected daily in pre-weighed sterile Eppendorf tubes, and fecal pellet weight was measured. Pellets were homogenized in 0.1% Proteose peptone by affixing them to a vortexer for 5 minutes. Homogenized stool was serially diluted in a 96-well plate and dilutions were plated on LB plates supplemented with kanamycin and MacConkey agar plates.

Mice were euthanized and tissue samples (Ileum, cecum, colon) were resected, weighed and resuspended in 0.1% proteose peptone. Tissue samples were homogenized using TissueLyser II (Qiagen) at 25 Hz for 90 seconds and serially diluted in proteose peptone. Dilutions were plated on LB-kanamycin and MacConkey agar plates and also inoculated into lactose broth and incubated at 37C aerobically for 16-20 hours. Additionally, glycerol stocks of homogenate were prepared (25% v/v). Colony-forming units were enumerated the following day.

### Barcode library preparation

gDNA was extracted from fecal and tissue samples using the MagAttract Power Microbiome DNA/RNA KF kit (Qiagen) and Kingfisher Flex (Thermo Fisher Scientific). Amplicon libraries were prepared using 50 ng of fecal gDNA for initial amplification of samples (BC amp Set 1 primers, Supplementary Table 1). 5 µl of this reaction was used in a subsequent round of amplification using BC amp set 3 primers to add Illumina adapters to the sequences followed by amplification using the Nextera XT DNA Library Preparation Kit (Illumina cat. FC-131-1096) to add 5’ and 3’ indices to the library. The bands corresponding to the amplicon were gel extracted and pooled together before sequencing.

### Quantitative PCR for assessing *E. coli* GI levels

Bacterial load in stool was quantified by qPCR analysis (SsoAdvancedSYBR Green Supermix, Bio-Rad) of microbial gDNA using the *E. coli uidA* gene as previously described.^81^ Bacterial abundance was determined using standard curves constructed with reference to cloned DNA corresponding to a short segment of the *uidA* gene that were amplified using conserved specific primers (qPCR Primer set 1). 25ng of microbial gDNA was used as an input template. Reaction conditions were 30 sec at 95°C, followed by 40 cycles of 1 sec at 95°C and 10 sec at 57.2°C.

### Barcode analysis

Amplicon sequencing was performed (BGIseq 2000 platform, PE 100). We quantified barcode frequencies from the raw sequencing reads with following steps:

1. Merged paired reads into a single longer reads (median read length 127) using FLASH1 with parameters minimum overlap is 10bp and maximum mismatch density is 0.25
2. Aligned merged reads to the amplicon region template sequence using bowtie22 with parameters set to very sensitive local alignment while disabling gaps (complete command included in 25-bowtie2-align-merged-pairs.run) in the reference sequence in order to eliminate reads with partial barcode integration
3. Counted successfully aligned reads using a custom python script (30-find-barcodes.py) that indexes each read with the aligned barcode sequence.
4. Generated day-by-day barcode frequency trajectories using a custom python script (40-create-barcode-trajectories.py) by normalizing barcode read counts to the total recovered barcode count at each sample.

### Cefepime dose-response curve assays

Overnight cultures of the strains were prepared and inoculated into a 96-well plate containing 2-fold dilutions of cefepime in LB media to a final OD_600_ of 0.001. Plates were incubated at 37°C with shaking (400 rpm in Infors HT plate shaker). Plates were read the next day using Biotek Epoch2 plate reader. Background subtracted data was obtained from Biotek Gen5 software after specifying blank wells. The minimum cefepime concentration where no bacterial growth was detected using optical density reads was designated as the MIC value (Background corrected OD_600_ < 0.01).

### Growth curve assays

Overnight cultures of the strains were prepared and inoculated into a 96-well plate containing LB to a final OD_600_ of 0.001. Plates were incubated in stacker device provided with Biotek Epoch2 machine and were read every 15 minutes for 24 hours. Growth curve data was obtained after background subtraction by Biotek Gen5 software.

### Cell invasion assay

Colorectal adenocarcinoma cell line HT-29 cells were grown in DMEM (Gibco cat. 11965092) (supplemented with 10% FBS and 1% Antibiotic-Antimycotic Gibco cat. 15240062) for 72 hours at 37°C until confluent. After washing once with pre-warmed 1X PBS, *E. coli* (1×10^5^ or 1×10^6^ CFU) were inoculated in fresh, pre-warmed DMEM media (supplemented with 10% FBS and further supplemented with chloramphenicol 25 µg/mL for plasmid-carrying cells and 50 mM arabinose for complemented strain) and added to 1×10^5^ HT-29 cells. After 4 hours of incubation at 37° C, the media was removed, and the cells were washed with ice cold PBS x 3. Pre-warmed DMEM supplemented with 100 µg/mL of gentamicin was then added to the HT-29 cells for 2 hours to kill any extracellular bacteria. HT-29 cells were then washed with ice cold PBS x 3 (washes were saved and plated). The cells were then lysed with 0.25% Trypsin-EDTA (Gibco 25200056) +0.5% Triton X 100 and the lysate was plated to enumerate the number of internalized bacteria.

### Cefepime time-kill assay

Overnight cultures of PbEc and IM6 were diluted 1:1000 and allowed to grow for 2 hours aerobically. The initial CFU of the culture was recorded using serial dilution and plating on LB agar. The cultures were then exposed to different concentrations of cefepime and every hour samples were obtained, washed twice with 1X PBS to remove residual antibiotic and serial dilutions and plating was used to assess CFU at different time points.

### Cefepime evolution assay

PbEc and IM6 (six replicates of each strain) were grown in LB with cefepime concentrations corresponding to 0.25X, 0.5X, 1X, 2X and 4X MIC. Based on growth (OD_600_>0.5), new MIC’s were calculated daily. Cultures were then diluted 1:100 into fresh media containing the aforementioned concentrations of cefepime. The experiment was carried out for 21 days. Two replicates of each strain were also simultaneously passaged in LB (without cefepime) for comparison.

### Colonocyte intracellular cefepime concentration assay

HT-29 cells were seeded in 6 well plates with 7.5 × 10^5^ cells per well and grown in DMEM media supplemented with 10% FBS. After 24 hours, the media was replaced with fresh media including 0.1, 1 and 5 µg/mL of cefepime, respectively. After 0.5, 2, 6 and 24 hours of incubation with compounds, cells were harvested using trypsin and washed in PBS. All the steps were carried out under ice cold conditions. Then each sample was diluted in PBS to achieve a final concentration of 1.5 × 10^6^ cells/mL. In parallel, untreated HT-19 cells were prepared with PBS to yield 1.5 × 10^6^ cells/mL (10 mL). All samples were snap-frozen in 15 mL conical tubes with liquid nitrogen and stored at -80°C for cefepime quantification by LC-MS/MS as described above. Total cefepime amounts were determined by multiplying the lysate concentration by the total lysate volume. The total number of cells contained in that lysate volume was then multiplied by the volume of an individual HT29 cell determined using a Scepter™ Handheld Automated Cell Counter (Millipore,Burlington, MA). Intracellular cefepime concentrations were then calculated by dividing the total lysate cefepime by the intracellular volume of cells in the lysate.

### LPS extraction and PAGE

LPS was extracted from the strains as previously described^82^. 10 µl of LPS extract was mixed with 10 µl 2X SDS dye and was run on 10% unstained polyacrylamide gel at 180 V for 30 minutes. Gel was stained using Pro-Q™ Emerald 300 Lipopolysaccharide Gel Stain Kit (ThermoFisher cat. P20495) and visualized using a BioRad imager.

### Statistical Analysis

GraphPad Prism v.9.2 was used for statistical analysis. All data sets were tested for normality (e.g. Shapiro-Wilk). Data sets with normal distribution were analyzed with parametric tests, such as standard student t-test or one-way ANOVA with Bonferroni post-test. For non-normal distributions, non-parametric tests, such as Mann Whitney U test or Kruskal Wallis with Dunn’s post-test, were applied.

## RESOURCE AVAILABILITY

### Lead contact

Further information and requests for resources and reagents should be directed to and will be fulfilled by the lead contacts Erdal Toprak and Andrew Koh

### Materials availability

Bacterial strains and mutants generated for this study will shared upon request

### Data and code availability

The sequencing data for will be deposited in the NCBI Sequence Read Archive.

Original code generated for this study will be deposited at github.

These processes are currently pending.

