## Supplementary Material for "Susceptible bacteria survive antibiotic treatment in the mammalian gastrointestinal tract without evolving resistance"

**This PDF file includes:**

Figures S1-S11

Table S1-S2

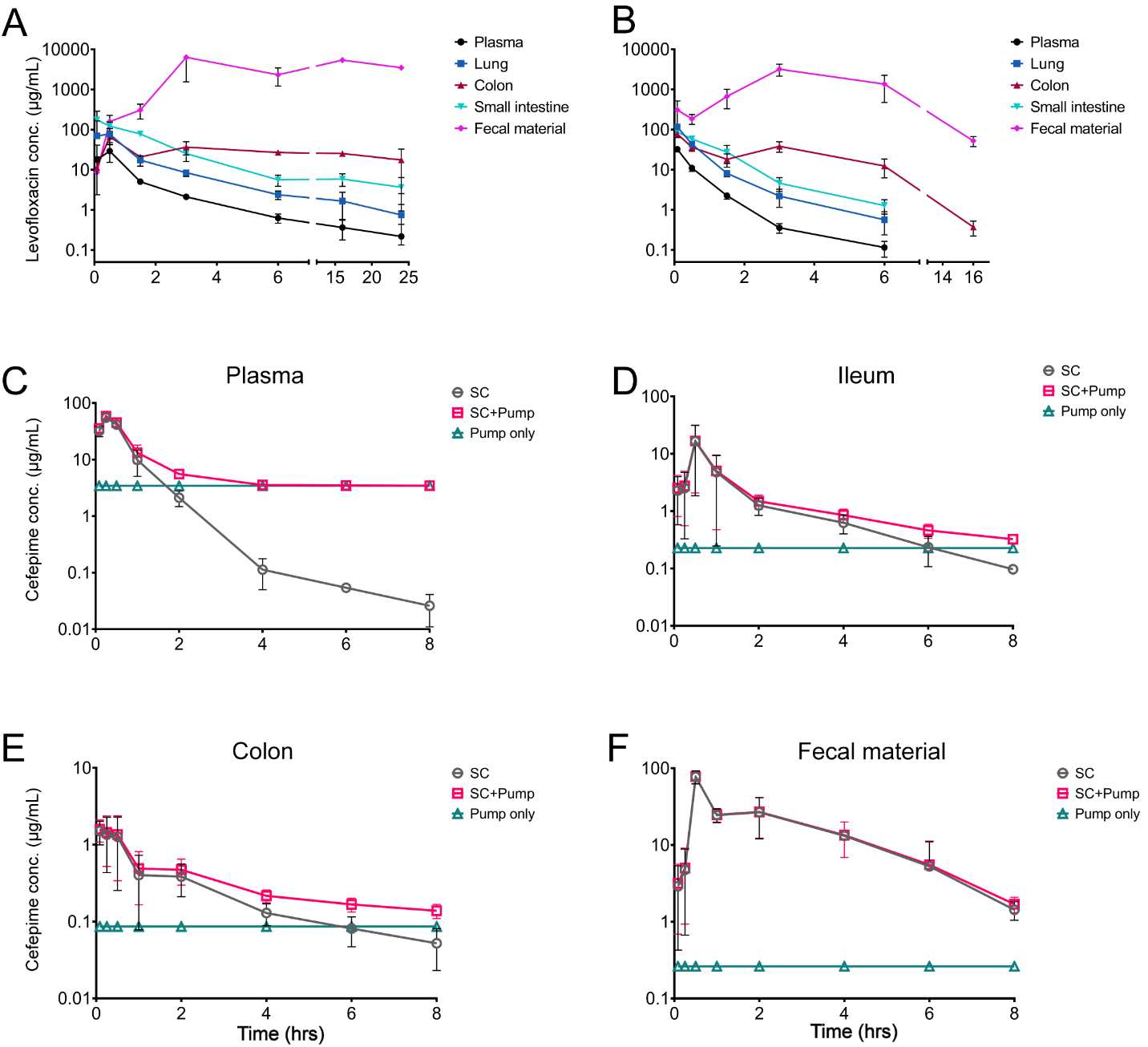

**Figure S1. Pharmacokinetics/pharmacodynamics measurements for levofloxacin and cefepime administered to conventional C57BL/6J mice.**

A. Concentration of levofloxacin in mouse plasma, lung, colon, ileum and fecal material over time after a single dose of levofloxacin administered through oral gavage (PO) (150 mg/kg) measured using Liquid-chromatography-Mass Spectrometry/ (LC-MS/MS.) n=3 mice per time point. Points represent the mean value per group ± SD.

**B.** Concentration of levofloxacin in mouse plasma, lung, colon, ileum and fecal material over time after a single dose of levofloxacin administered through IV (75 mg/kg) measured using LC-MS/MS. n=3 mice per time point. Points represent the mean value per group ± SD.

**C.** **Plasma, D. Ileum, E. Colon, F. Fecal material** concentrations of free cefepime in mice (C57BL/6J, Jackson, female, 6-8 weeks) after single subcutaneous (SC) dose (50 mg/kg dose x 1) or after continuous cefepime dosing (iPrecio SM310R pump, 20 mg/mL, 4 µl/hr) measured using LC-MS/MS (Shimadzu Prominence HPLC coupled to a Sciex 4000 QTRAP(r) mass spectrometer). n=3 mice per group. SC + Pump values were calculated by mathematical addition of SC concentrations to pump concentrations. Points represent the mean value per group ± SD.

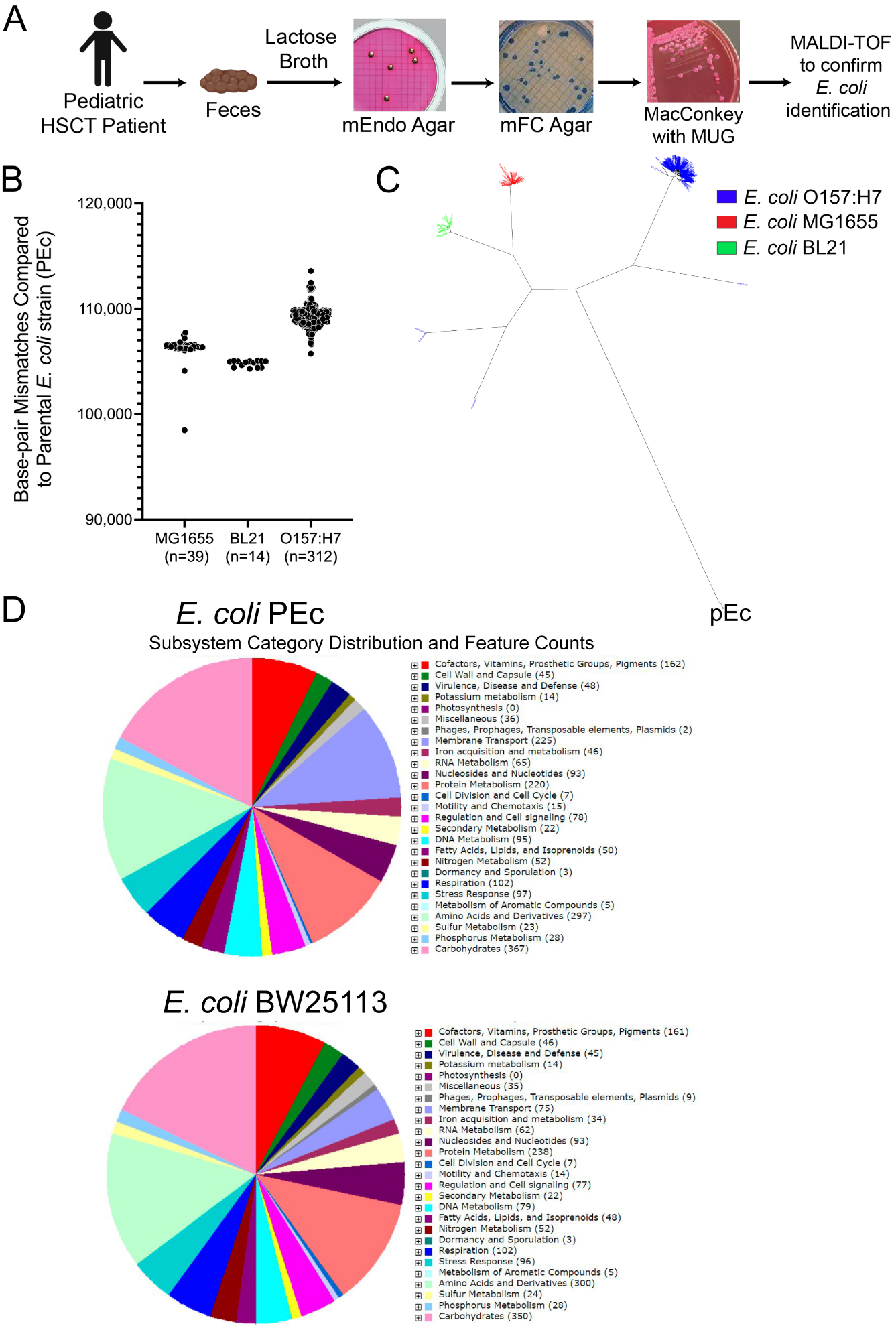

**Figure S2. Isolation and characterization of a pan-sensitive clinical *E. coli* strain (PEc)**

**A.** Overview of *E. coli* isolation protocol implemented from patient stool sample prior to exposure to antibiotics or chemotherapy. After initial enrichment in lactose broth, successive plating on selective agar: mEndo followed by mFC agar and finally MacConkey Agar with MUG. Colonies appearing fluorescent on this media were confirmed as *E. coli* (designated PEc) using MALDI-TOF (Bruker MALDI Biotyper).

**B.** Comparison of base-pair mismatches (i.e., single nucleotide polymorphisms, SNPs) found in PEc and commonly used *E. coli* lab strains: MG1655, BL21 and O157:H7. Whole genome sequencing on PEc was performed (HiSeq platform, PE150, Seqcenter, PA,). MUMmer 4^1^ was used to compare the genome assemblies. The mutation rate between two genomes was calculated by the number of SNPs divided by the alignment length.

**C.** Dendogram depicting clustering of PEc compared to commonly used lab strains from **B.** The mutation rate was used to calculate the Jukes-Cantor distances. The neighbor-joining tree was generated based on the distances using R-packages ape^2^ and ggtree ^3^.

**D.** RAST (Rapid Annotation using Subsystem Technology)^4^ analysis of *Escherichia coli* strains PEc and BW25113. Illumina sequencing reads of PEc (PE-150) were assembled into contigs using CLC Genomics Workbench Denovo Assembly tool. Contigs were submitted to RAST for gene annotation and prediction. Pie chart obtained from RAST output showing sub-system category distribution of genes found in PEc compared to *E. coli* BW 25113 (NCBI Accession no. CP009273.1).

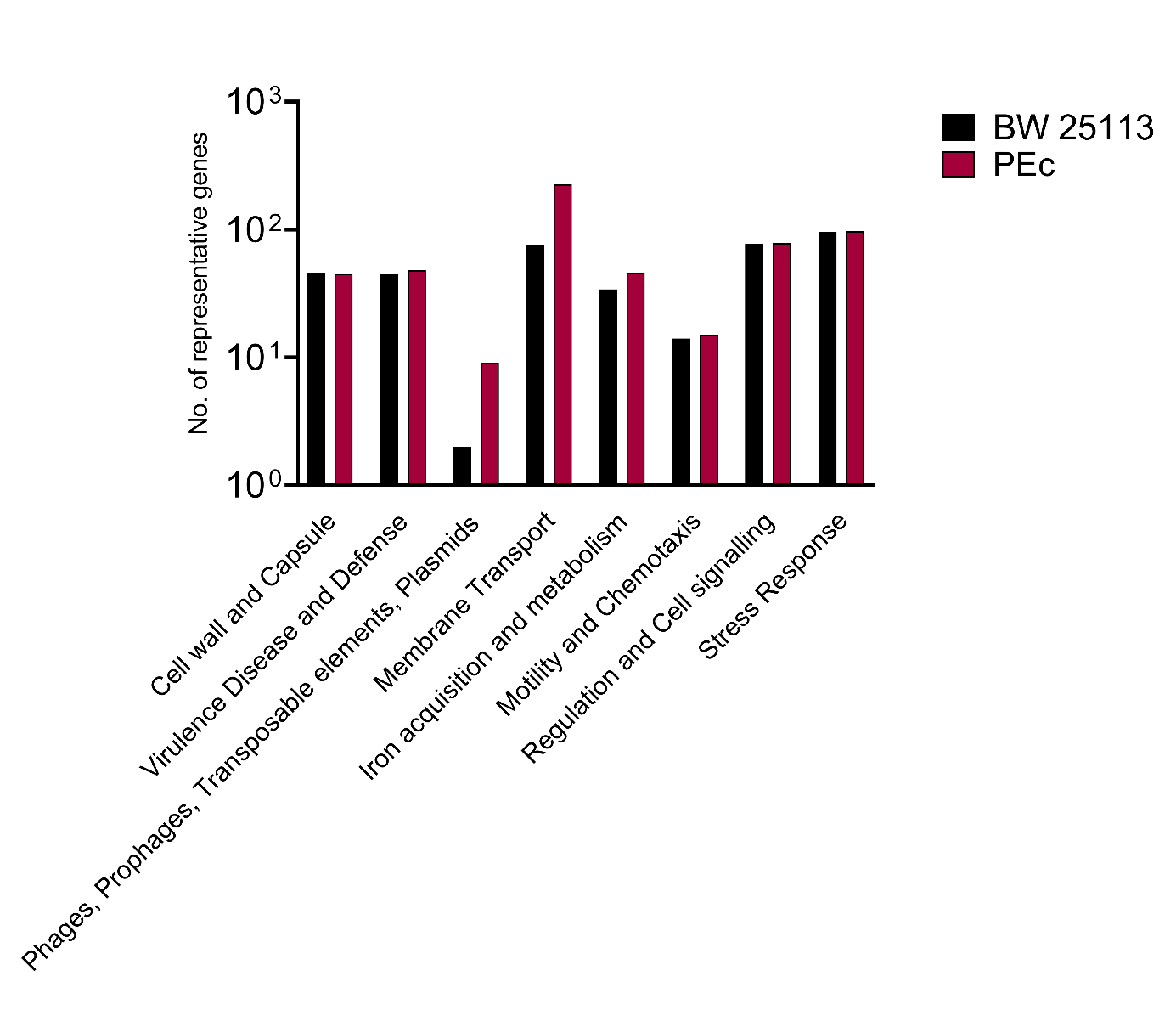

**Figure S3. Comparison of RAST-identified gene sub-groups and classifications of *E. coli* PEc strain and *E. coli* BW25113.** Select gene sub-groups and classifications compared between *E. coli* PEc and *E. coli* BW25113.

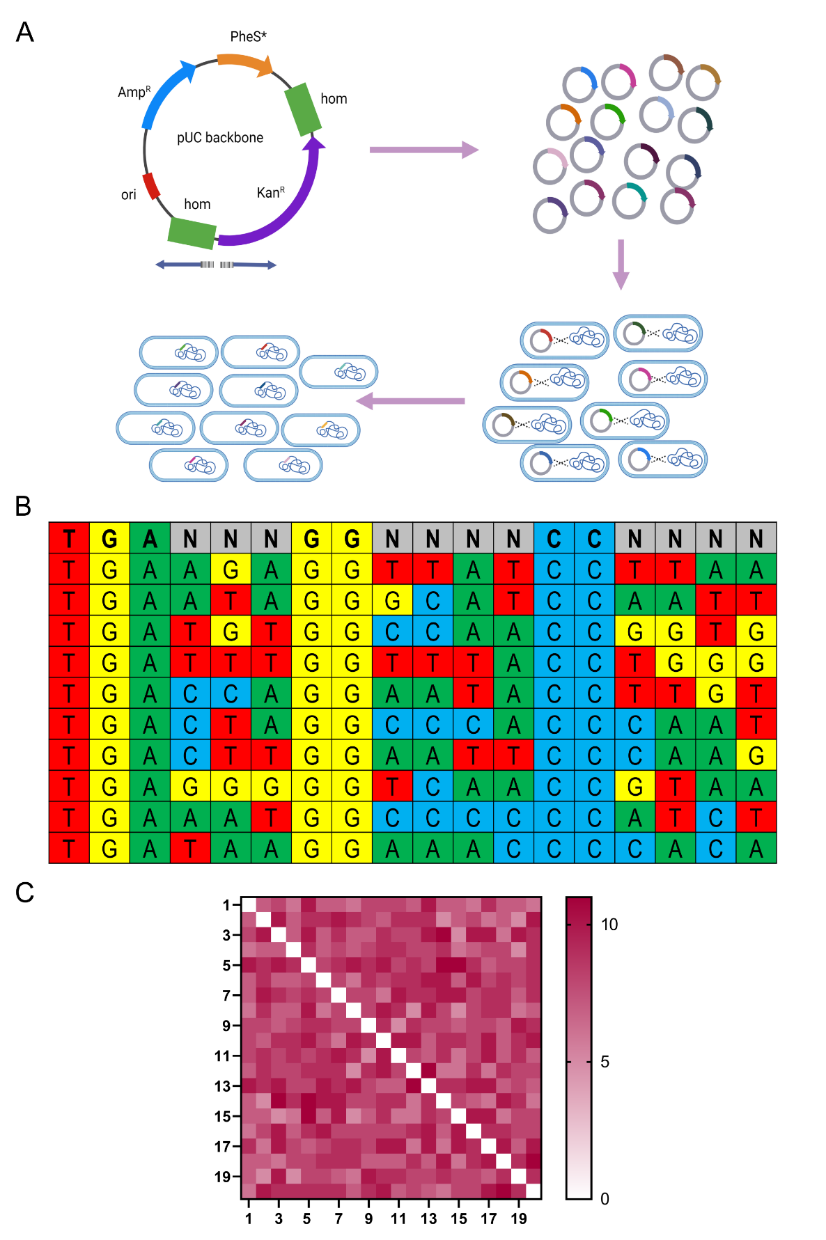

**Figure S4. Genome barcoding protocol and barcode analysis overview.**

**A.** Overview of barcode library construction protocol. Plasmid bearing pUC origin of replication was engineered to carry Ampicillin resistance gene on its backbone, a counter selective gene PheS*(Phenylalanine tRNA synthetase T251A/ A294G)^5^, and two homology regions flanking a Kanamycin resistance cassette for recombination into the genome. The plasmid was linearized using primers with randomized barcode sequences with NNN base pairs on their 5’ end and then ligated and transformed into NEB10β cells to create the barcoded plasmid library. This library was then transformed into PEc cells induced for lambda red recombineering (with pKD-PC, a pKD46 derivative engineered to express chloramphenicol resistance and the counterselective gene *pheS**). After transformation, cells were passaged in M9 + kanamycin supplemented with p-cl-phenylalanine to enforce selection for integrants while simultaneously counter-selecting against both plasmid backbones until no ampicillin-resistant or chloramphenicol-resistant clones remained.

**B.** Diversity of the integrated barcodes as compared to the initial library. Genomic DNA isolated from PbEc was amplified (BC amp primers set 1 and 2, in Supplementary Table 1). Amplicon sequencing was performed by Azenta Inc. (Amplicon-EZ service). A custom python script was used to identify the barcodes in the amplicons. The figure depicts an alignment of 10 barcodes from the analysis to the randomized barcode sequence (first row) displaying the diversity of the library.

**C.** Heatmap showing the genetic distance (in units of nucleotide counts) between the top 20 most frequent barcodes identified in the above round of sequencing.

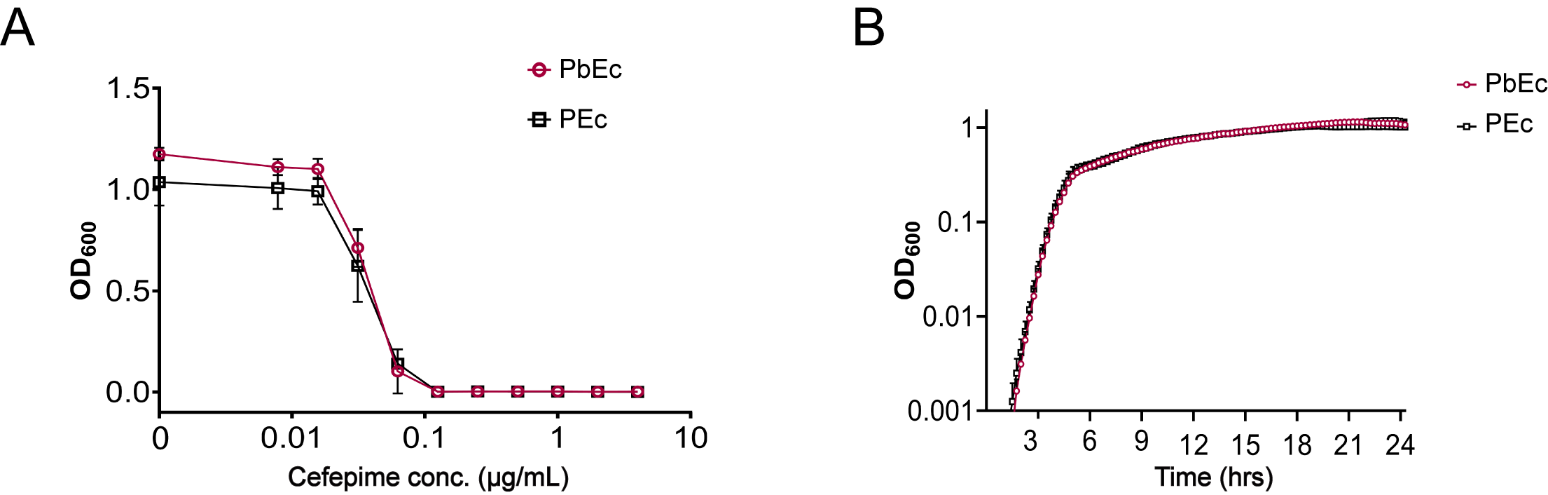

**Figure S5. Phenotypic comparison of PEc to PbEc.**

A. **Cefepime susceptibility of PEc remains unchanged after barcoding.**

Cultures of PEc and PbEc (barcoded PEc) were grown overnight in LB and added to a 96-well plate containing serial dilutions of cefepime (2-fold) to a starting OD_600_ of 0.001. Plates were incubated (Infors HT at 400 rpm, 37°C) overnight and growth assessed the following OD_600_ measurement. Points represent mean ± SD. 3 technical replicates were performed for each experiment. 3 biological experiments were performed. Cefepime MIC for both PEc and PbEc are ~0.1 μg/mL.

B. **Barcoding has no detectable effects on growth phenotype of PEc.**

Cultures of PEc and PbEc were grown overnight in LB and added to a 96-well plate containing LB to a starting OD_600_ of 0.001. Plates were incubated and OD_600_ was measured every 15 minutes. Points represent mean ± SD. 3 technical replicates were performed for each experiment. 2 biological experiments were performed. Doubling time for PEc (21.65 mins) vs PbEc (20.48 mins).

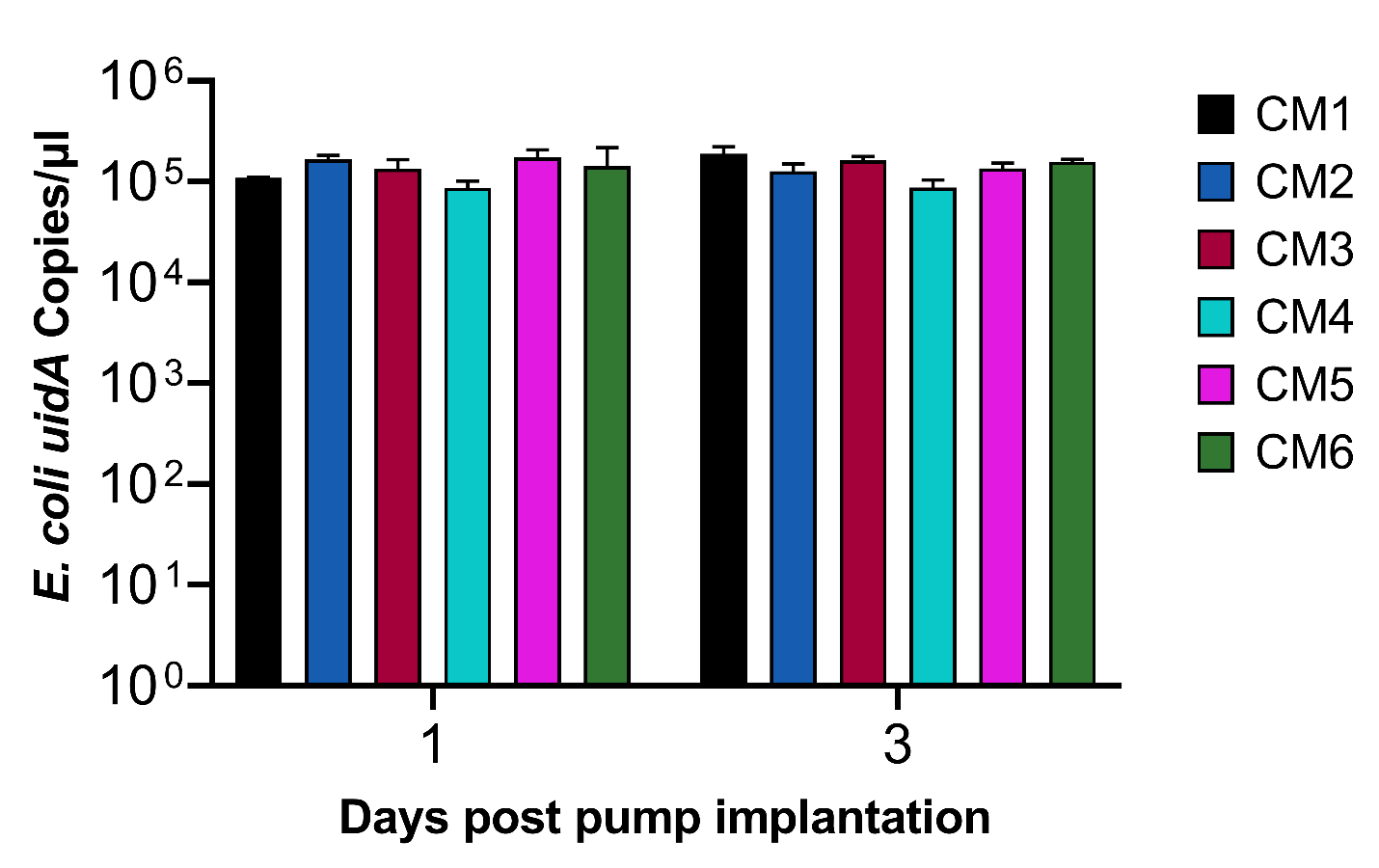

**Figure S6. *E. coli* GI colonization levels of control mice.** *E. coli* GI levels as determined by qPCR of fecal gDNA of *E. coli* specific *uidA* gene. Fecal samples collected from control mice (CM1-CM6) on Days 1 and 3. Bars represent mean ± SD for each mouse. 2 technical replicates performed for each mouse sample

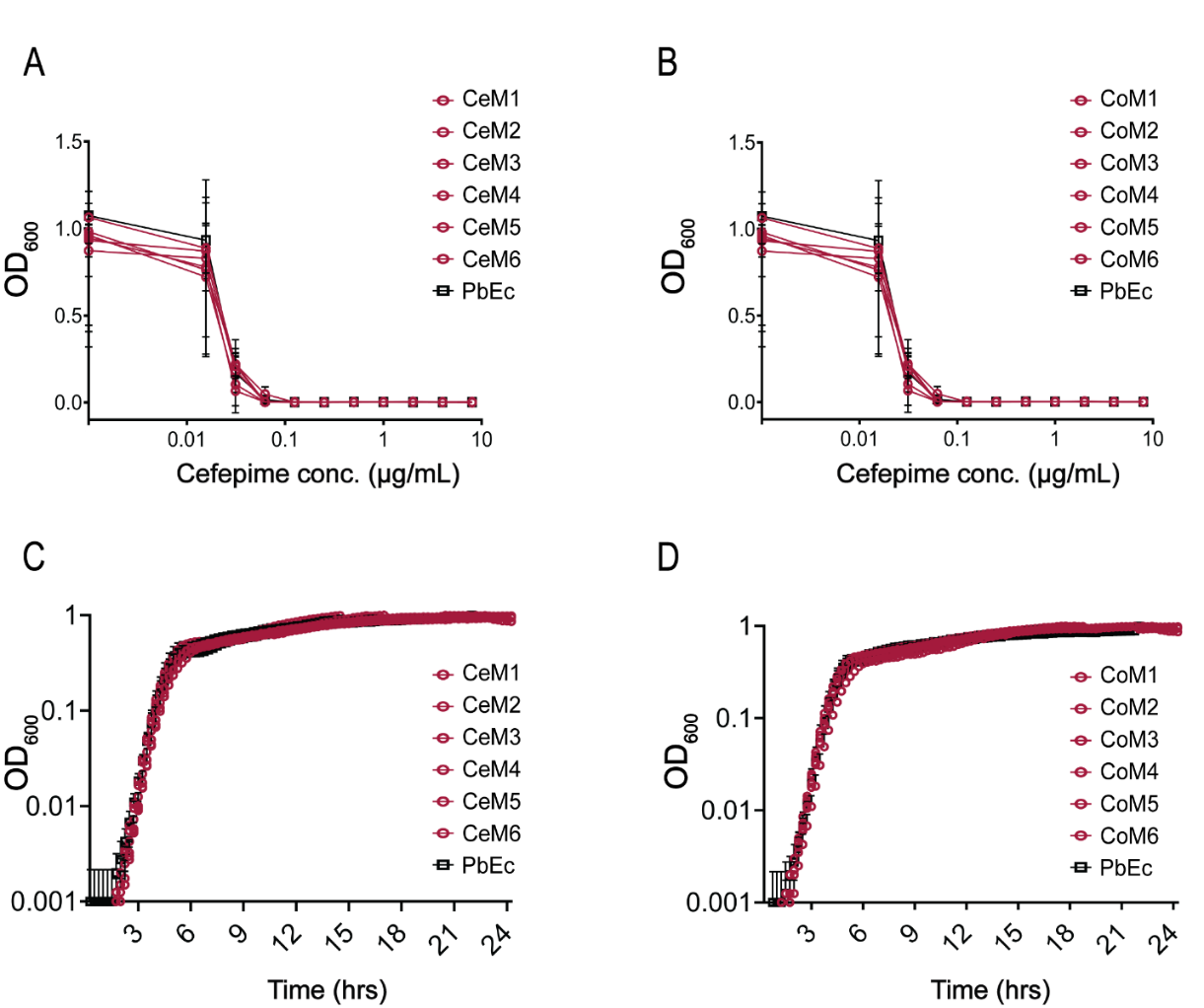

**Figure S7. Phenotypic characterization of PbEc *E. coli* isolates recovered from GI tissue samples from cefepime-treated mice.**

**A-B. PbEc cultures recovered from tissue of cefepime treated mice are not resistant to cefepime.** Cecal *E. coli* isolates **(A)**, colon *E. coli* isolates **(B)**, and PbEc were grown overnight in LB and added to a 96-well plate containing serial dilutions of cefepime (2-fold) to a starting OD_600_ of 0.001. Plates were incubated (Infors HT at 400 rpm, 37°C) overnight and growth assessed the following OD_600_ measurement. Points represent mean ± SD. Three replicates performed for mixed cultures derived from each tissue. No cultures grew at 0.1 µg/mL or higher concentrations (data not shown)

**C-D.** **PbEc isolates recovered from cefepime-treated mice do not exhibit growth differences compared to the parental PbEc strain**

Cecal *E. coli* isolates **(C)**, colon *E. coli* isolates **(D)**, and parental PbEc were grown overnight in plain LB and added to a 96-well plate containing plain LB to a starting OD_600_ of 0.001. Plates were incubated and OD_600_ was measured every 15 minutes. Points represent mean ± SD. Three replicates performed for each isolate.

CeM1, *E. coli* isolate recovered from cecum of mouse 1 of cefepime-treated group. CeM2, *E. coli* isolate recovered from cecum of mouse 2 of cefepime-treated group, etc.

CoM1, *E. coli* isolate recovered from colon of mouse 1 of cefepime-treated group. CoM2, *E. coli* isolate recovered from colon of mouse 2 of cefepime-treated group, etc.

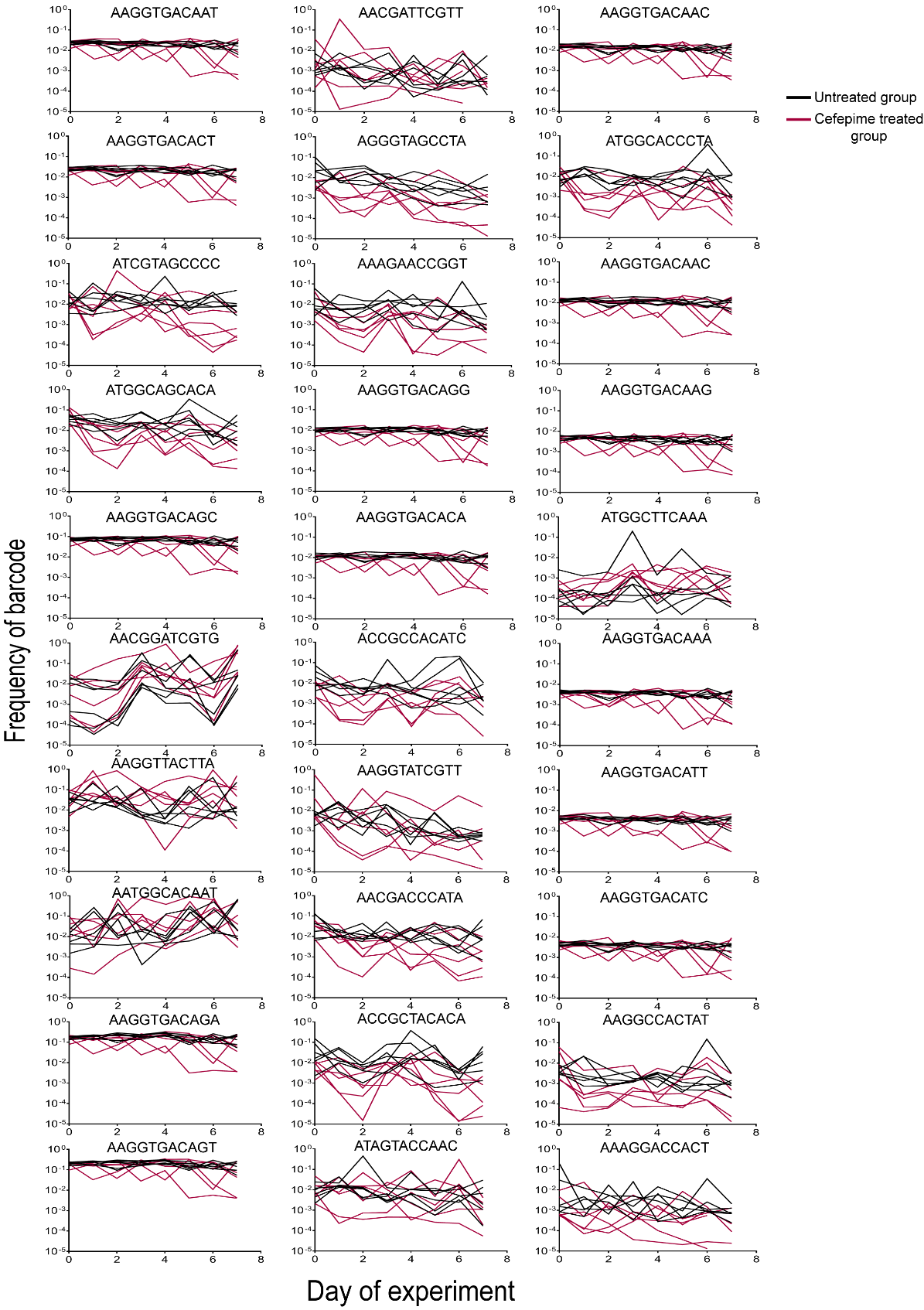

**Figure S8. Frequency of the 30 most commonly observed barcodes of PbEc in germ-free mice treated with or without cefepime.** Fecal gDNA was extracted from daily stool samples from individual mice. Barcoded genomic regions were amplified (from daily fecal gDNA samples) for amplicon sequencing. **Red** lines indicate cefepime-treated mice, **black** lines are control (untreated mice).

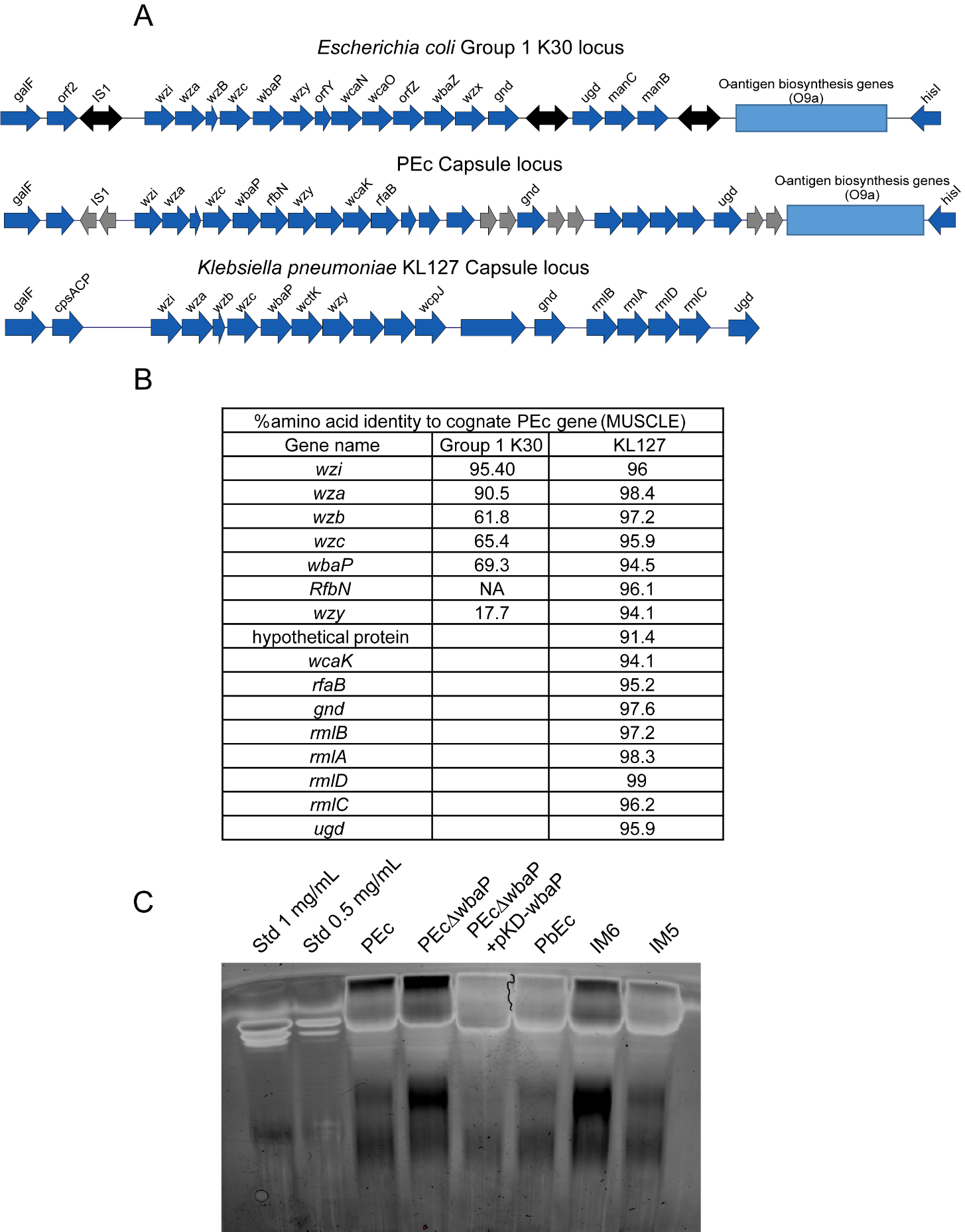

**Figure S9. PEc capsule locus and lipopolysaccharide (LPS) analysis.**

**A.** Capsule locus structures of *E. coli* Group 1 capsule, PEc and *Klebsiell*a KL 127 capsule locus type. Comparison of capsule loci structures of canonical *E. coli* Group 1 K30 capsule (adapted from^6^), PEc capsule locus and *Klebsiella pneumoniae* KL 127 capsule locus. PEc capsule locus was identified as Group 1 *E. coli* capsule based on the presence of the *wzi* gene and the absence of the colanic acid biosynthesis genes. Similarity to KL 127 capsule locus was confirmed after using BLASTn to identify unknown genes in PEc locus.

**B.** Table comparing protein sequence identity (using MUSCLE) of corresponding genes in *E. coli* Group 1 and *Klebsiella* KL 127 to PEc genes (shown as percentage amino acid identity of genes in capsule locus compared to *E. coli* PEc)

**C.** LPS gel image of PEc, PEc-ΔwbaP, PbEc, IM6 and IM5**.** LPS was extracted from each sample followed by PAGE using a 10% mini-Protean TGX stain-free gel 180 V, 30 min) followed by staining using Pro-Q™ Emerald 300 Lipopolysaccharide Gel Stain Kit).

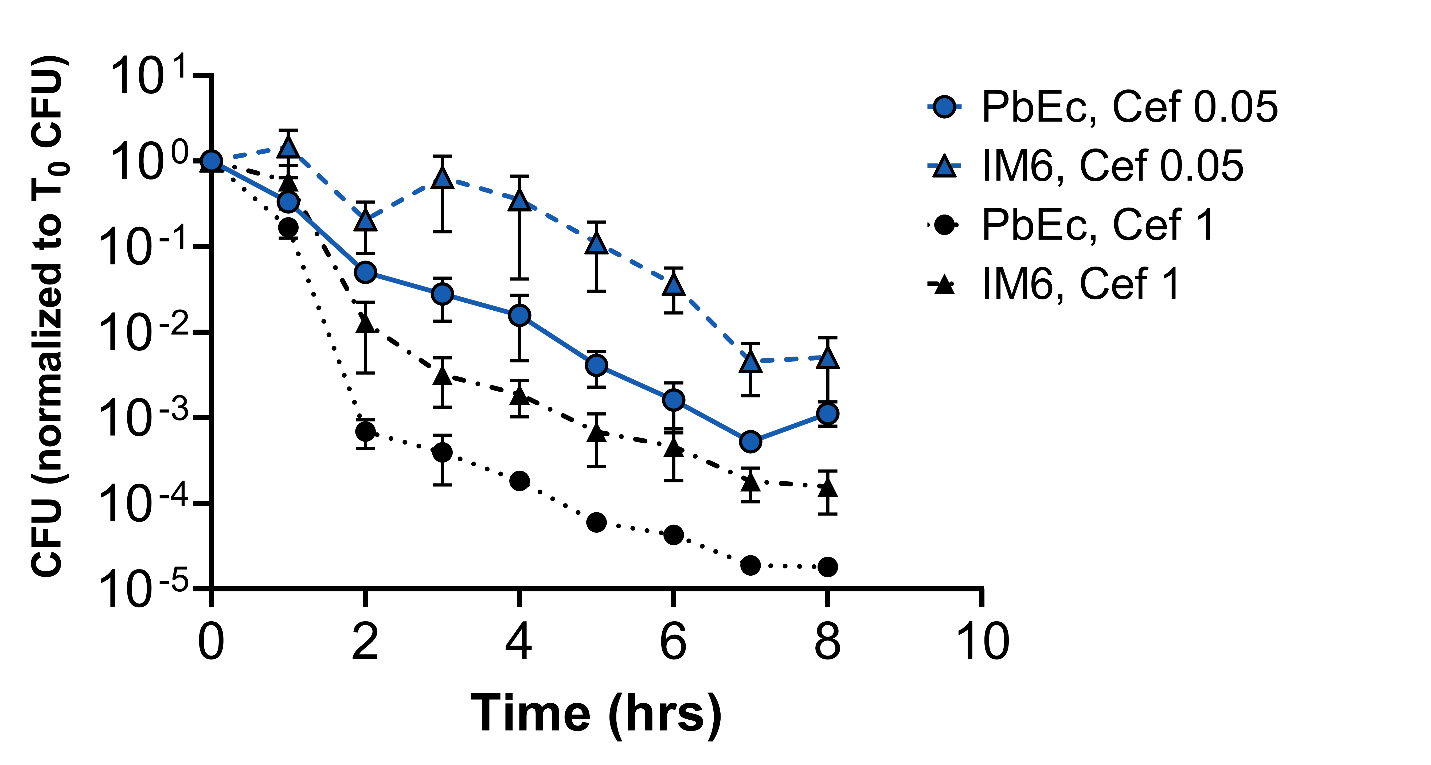

**Figure S10. Time kill assay for parental PbEc library and IM6 isolate at two different cefepime concentrations.** Overnight cultures of PbEc and IM6 (n = 6 replicates) were diluted 1:1000 in LB and re-incubated at 37°C with shaking until at log phase growth. Initial timepoint (T_0_) *E. coli* quantification was performed (by enumeration of cultured CFD) and then cefepime (1 µg/mL or 0.05 µg/mL final concentrations) was added to each of the cultures and incubated at 37°C with shaking. Every hour, *E. coli* growth/survival was assessed by enumeration of cultured CFU. Points represent the mean ± SEM.

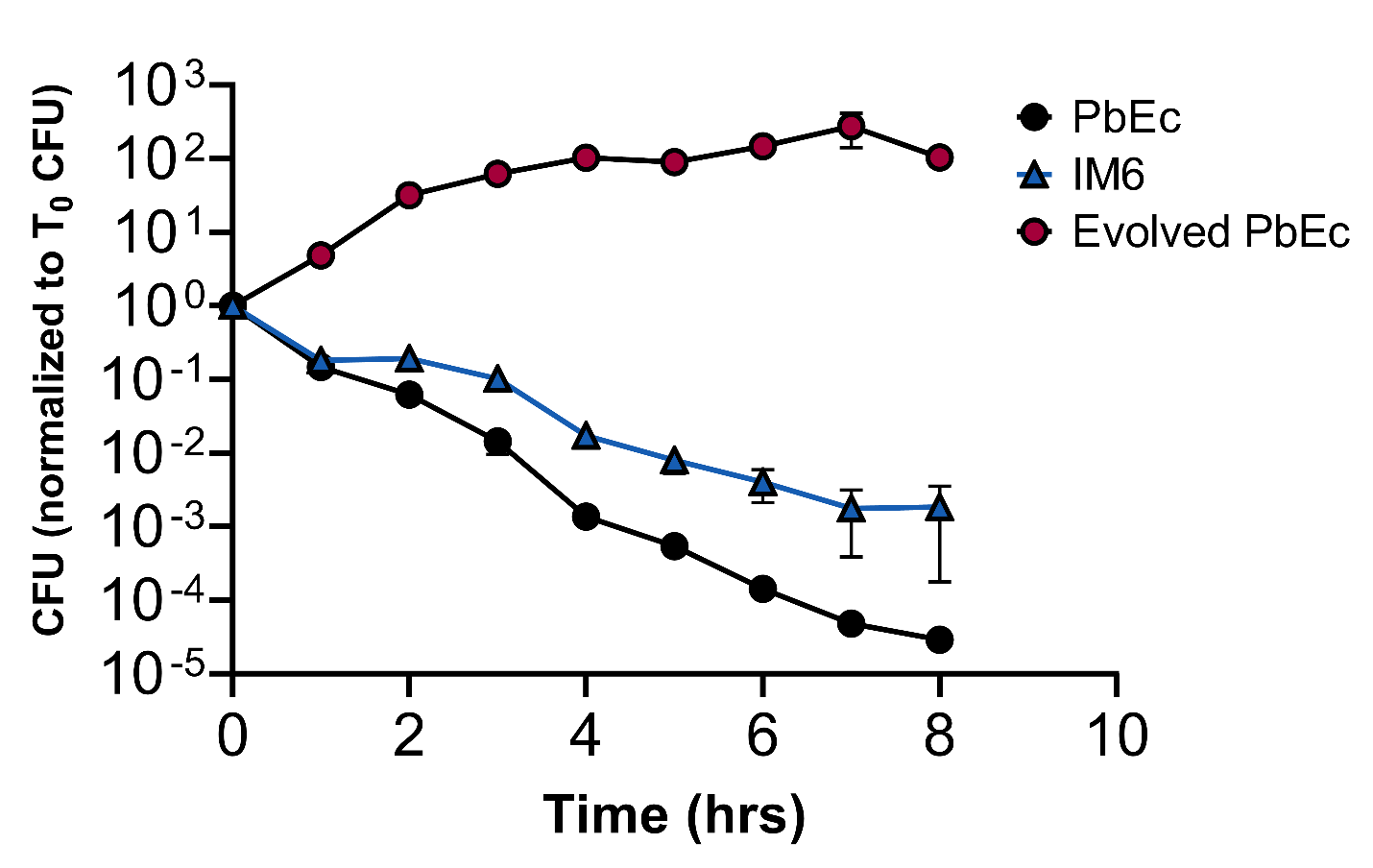

**Figure S11. Time kill assay for parental PbEc, IM6 isolate and cefepime-evolved PbEc (after 21 days).**

Time-kill assay performed in the presence of 1 µg/mL cefepime as described in **Fig. S10** for strains PbEc, IM6, and a laboratory evolved cefepime-resistant strain PbEc obtained from evolution experiment (**Fig. 5E**) n=6 replicates per strain. Points represent the mean ± SEM.

**Supplementary Table 1**

| Primer set name | Primer set name | Sequence |
| --- | --- | --- |
| Primers for creating barcoded library | | |
| BC set 1 | Homology 1 primers | GTGAAAACCTCTGACACATGGTTCGATCATGAAAAACTGTAAATAACG |
|  |  | AGTAAATAAATATGCTGTGCGCGAAC |
| BC set 2 | Kanamycin resistance gene primers | CCGATCTGTTTAAACCTAG |
|  |  | TTGCAAAGGCATCATTTGCCATATGTGGGAGTATAAGACG |
| BC set 3 | Homology 2 primers | TCTTATACTCCCACATATGGCAAATGATGCCTTTGC |
|  |  | TATCCGTTGCTGAATCTGAAATGTCAATGGGTGGTTTTTG |
| BC set 4 | pUC backbone amplification primers | AAAAACCACCCATTGACATTTCAGATTCAGCAACGGATAC |
|  |  | ACAGTTTTTCATGATCGAACCATGTGTCAGAGGTTTTC |
| BC set 5 | Primers to linearize plasmid to add PheS | ATTACACCTGATGAGATCTCATGACCAAAATCCCTT |
|  |  | TTCAGTGAACTGTGGTTATCAAAAAGGATCTTCACC |
| BC set 6 | primers amplifying PheS | CTCATCAGGTGTAATTGTC |
|  |  | CCACAGTTCACTGAATTTC |
| BC set 7 | primers adding spe promoter to PheS* | TAGCTCAGTCCTAGGTATAATACTAGTATGTCACATCTCGCAGAACTG |
|  |  | TATTATACCTAGGACTGAGCTAGCTGTCAAGGTTTTCCTCATTGTGTCAGTG |
| BC set 8, 9 | primers adding 2 SNPs to PheS to generate PheS* | GGCTTCGGGATGGGGATGGAGCG |
|  |  | TTCCGCAAACGGGAAGTAGGAAGGACG |
|  |  | TTTACCGAACCTTCTGCAGAAGTG |
|  |  | AAGCCGAAACCAGAGTAAACTTC |
| BC set 10 | Primers to linearize plasmid with barcode overhangs | /5Phos/GTNNNNGGNNNNCCNNNTCAGCTAGTAAATAAATATGCTGTGCGCGAAC |
|  |  | /5Phos/TGNNNTCAGNNNTTNNNATGACGCCGATCTGTTTAAACCTAG |
| pKD set 1 | Primers to linearize pKD plasmid for CamR cassette | AATAAGCGGATGAATGGCAGAAACTCATGAGCGGATACATATTTG |
|  |  | AAAGTTGGAACCTCTTACGTGCAAAAGGATCTAGGTGAAGATCC |
| pKD set 2 | Primers to amplify CamR gene | GCACGTAAGAGGTTCCAAC |
|  |  | TTTCTGCCATTCATCCGCTT |
| pKD set 3 | Primers to linearize pKD plasmid for PheS gene | TTCAGTGAACTGTGGATGTAACGGTGAACAGTTG |
|  |  | ATTACACCTGATGAGAATCAAAGGGAAAACTGTCC |
| Primers for creating wbaP deletion mutant | | |
| ΔwbaP set 1 | Homology 1 primers | gtgaaaacctctgacacatgATGGTGCCTCTAACTAGAAAC |
|  |  | gaagttcctatactttctagagaataggaacttcAAATAAAAAAAATCGAAACATATAATGGAATAAC |
| ΔwbaP set 2 | Kanamycin resistance gene primers | gaagttcctattctctagaaagtataggaacttcATTGATAGTCTGATCGGTCAACGTATAATCGAGTC |
|  |  | gaagttcctatactttctagagaataggaacttcCGCGCAGCGTCGCATCAG |
| ΔwbaP set 3 | Homology 2 primers | gaagttcctattctctagaaagtataggaacttcTCTTGATACTTGGTATGTCAAAAATTG |
|  |  | gcatatgtgggagtataagaTTAATACGCGCCATCTTTTTTTAAG |
| ΔwbaP set 4 | Primers to linearize pUC plasmid to add deletion construct | TCTTATACTCCCACATATGC |
|  |  | CATGTGTCAGAGGTTTTC |
| Primers for creating wbaP complementation plasmid | | |
| wbaP complement set 1 | Primers to amplify arabinose inducible machinery from pKD | cgtcagatatGCACGTAAGAGGTTCCAAC |
|  |  | gaggcaccatTTTTTATAACCTCCTTAGAGCTC |
| wbaP complement set 2 | wbaP gene amplification primers | gttataaaaaATGGTGCCTCTAACTAGAAAC |
|  |  | tgtatcagtcTTAATACGCGCCATCTTTTTTTAAG |
| wbaP complement set 3 | Primers to amplify ori from pUC plasmid | cgcgtattaaGACTGATACAATCGATTTCTG |
|  |  | tcttacgtgcATATCTGACGATTGGACTTC |
| Primers for amplifying Barcode library | | |
| BC amp set 1 | Primers to create integrated barcode amplicons | TACAATTGCGACTTTTCTGC |
|  |  | GTTTGCAAAAGCTAGGACTC |
| BC amp set 2 | Primers to add Genewiz sequences to integrant amplicons | ACACTCTTTCCCTACACGACGCTCTTCCGATCTTACAATTGCGACTTTTCTGC |
|  |  | GACTGGAGTTCAGACGTGTGCTCTTCCGATCTGTTTGCAAAAGCTAGGACTC |
| BC amp set 3 | Primers to add Illumina sequences to integrant amplicons | TCGTCGGCAGCGTCAGATGTGTATAAGAGACAGNNNNNNTACAATTGCGACTTTTCTGC |
|  |  | GTCTCGTGGGCTCGGAGATGTGTATAAGAGACAGNNNNNNGTTTGCAAAAGCTAGGACTC |
| Primers for *E. coli uidA* qPCR | | |
| qPCR set 1 | Primers for E. coli uidA qPCR | CAATGGTGATGTCAGCGTT |
|  |  | ACACTCTGTCCGGCTTTTG |
|  | Probe for qPCR | 6FAM-TTGCAACTGGACAAGGCACCAGC-BBQ |

**Supplementary Table 1. Oligonucleotide primers used for this study.**

**Supplementary Table 2**

| PbEc 1 | | PbEc 2 | |
| --- | --- | --- | --- |
| EFFECT | PRODUCT | EFFECT | PRODUCT |
| missense_variant Val281Gly | tRNA-5-carboxymethylaminomethyl-2- thiouridine(34) synthesis protein MnmE | stop_gained Trp124* | Outer membrane porin OmpC |
| stop_gained Tyr35* | Outer membrane porin OmpC | synonymous_variant Ser87Ser | hypothetical protein |
| stop_gained Gln34* | Adenosine (5')-pentaphospho-(5'')-adenosine pyrophosphohydrolase | missense_variant Pro37Gln | HTH-type transcriptional repressor ComR |
| missense_variant Val838Ala | Protein RhsD | missense_variant Leu30Arg | Copper sensory histidine kinase CpxA |
| missense_variant Ala257Val | Cell division protein FtsI [Peptidoglycan synthetase] wbaP | missense_variant Val313Met | Cell division protein FtsI [Peptidoglycan synthetase] (EC 2.4.1.129) |
| frameshift_variant Phe32fs | Undecaprenyl-phosphate galactosephosphotransferase | frameshift_variant Phe32fs | Undecaprenyl-phosphate galactosephosphotransferase wbaP |
| missense_variant Gly290Arg | Multidrug efflux system AcrAB-TolC, inner-membrane proton/drug antiporter AcrB (RND type) | missense_variant Val127Gly | Multidrug efflux system AcrAB-TolC, inner-membrane proton/drug antiporter AcrB (RND type) |
| Promoter mutation | DNA-directed RNA polymerase beta' subunit RpoB | Promoter mutation | AcrR Transcriptional regulator of acrAB operon |
| Promoter mutation | MarR gene | missense_variant Arg77Cys | Multiple antibiotic resistance protein MarR |
|  | | frameshift_variant Thr412fs | ATP synthase beta chain |
|  |  | stop_gained Gln25* | ATP synthase epsilon chain |
| IM6 1 | | **IM6 2** | |
| EFFECT | PRODUCT | EFFECT | PRODUCT |
| stop_gained Gln171* | Outer membrane porin OmpC | missense_variant Val281Gly | tRNA-5-carboxymethylaminomethyl-2- thiouridine(34) synthesis protein MnmE |
| synonymous_variant Ser87Ser | hypothetical protein | stop_gained Tyr35* | Outer membrane porin OmpC |
| missense_variant Gly92Ser | Copper sensory histidine kinase CpxA | missense_variant Tyr141Asn | Copper sensory histidine kinase CpxA |
| stop_gained Gln240* | Fructose repressor FruR, LacI family | stop_gained p.Gln34* | Adenosine (5')-pentaphospho-(5'')-adenosine pyrophosphohydrolase |
| missense_variant Leu376Phe | Cell division protein FtsI [Peptidoglycan synthetase] | missense_variant Ala257Val | Cell division protein FtsI [Peptidoglycan synthetase] |
| missense_variant Ala498Val | Cell division protein FtsI [Peptidoglycan synthetase] wbaP | frameshift_variant Phe32fs | Undecaprenyl-phosphate galactosephosphotransferase wbaP |
| frameshift_variant Phe32fs | Undecaprenyl-phosphate galactosephosphotransferase | missense_variant Ile7Asn | IncF plasmid conjugative transfer pilus assembly protein TraF |
| missense_variant Gln213Leu | Multidrug efflux system AcrAB-TolC, inner-membrane proton/drug antiporter AcrB (RND type) | missense_variant Pro389Ser | Osmolarity sensory histidine kinase EnvZ |
| missense_variant Gln1326Leu | DNA-directed RNA polymerase beta' subunit RpoB | missense_variant Gly290Arg | Multidrug efflux system AcrAB-TolC, inner-membrane proton/drug antiporter AcrB (RND type) |
| missense_variant Asn266His | DNA-directed RNA polymerase beta' subunit RpoB | Promoter mutation | RpoB |
| frameshift_variant Lys62fs | Multiple antibiotic resistance protein MarR |  |  |
| stop_gained Gln10* | ATP synthase F0 sector subunit b |  |  |

**Supplementary Table 2. Comparison of mutations found in PbEc and IM6 after 21 days of cefepime resistance evolution.**

gDNA was isolated from Day 21 cultures of two PbEc evolved strains and two IM6 evolved strains. gDNA was submitted sequencing (Illumina, PE 150). Reads obtained were mapped to contigs obtained previously (**Fig. S2**) using the Snippy tool on usegalaxy.org. Mutations in common genes are highlighted in gray.
